# DSIF factor Spt5 coordinates transcription, maturation and exoribonucleolysis of RNA polymerase II transcripts

**DOI:** 10.1101/2023.10.16.562519

**Authors:** Krzysztof Kuś, Loic Carrique, Tea Kecman, Marjorie Fournier, Sarah Sayed Hassanein, Ebru Aydin, Cornelia Kilchert, Jonathan M. Grimes, Lidia Vasiljeva

## Abstract

The timely termination of RNA polymerase II (Pol II) transcription is critical for polymerase recycling and preventing interference with the expression of the neighbouring genes. Termination of Pol II transcription involves exoribonucleolytic decay of the nascent RNA by 5’-3’ exonuclease Xrn2. Xrn2 attacks the 5’-PO_4_-end and executes degradation of the nascent RNA generated by the endonucleolytic cleavage at the poly(A) site which eventually leads to Pol II release from DNA. However, the molecular details of when and how during this process Xrn2 interacts with the Pol II elongation complex to mediate its dissociation from DNA is not understood. Here, we demonstrate that Xrn2 interacts with Pol II and Spt5, a conserved transcription factor that controls Pol II processivity and pausing. Importantly, Xrn2 activity is stimulated by Spt5 *in vitro* and Spt5-depleted cells show defective transcription termination. Our results support a model where Xrn2 first forms a stable complex with the elongating Pol II to acquire its full activity in degrading nascent RNA. Spt5 also promotes premature termination attenuating the expression of non-coding transcripts. By contrast, Spt5 depletion leads to Pol II retention at promoters of protein-coding genes. Pol IIs that transcribe into a gene body in the absence of Spt5 exhibit severely reduced elongation rates and defective pre-mRNA processing. We propose that Spt5 plays a major role in the production of functional mRNA by directly stimulating the activity of the RNA enzymes and preventing entry of Pol II complexes not configured to support transcription and pre-mRNA processing into elongation.

## INTRODUCTION

DNA-dependent RNA polymerase II (Pol II) is responsible for the transcription of protein-coding (mRNA) and non-coding RNA (ncRNA) in eukaryotic cells. ncRNAs represent a diverse class of transcripts including stable house-keeping ncRNAs such as small nuclear RNAs, telomerase RNA and unstable ncRNAs such as enhancer (e)RNAs or promoter upstream transcripts derived from the bi-directional promoters^1^. Both classes of Pol II transcription units (TU) undergo maturation/processing to become functional molecules. RNA-processing of the nascent RNA takes place throughout the Pol II transcription cycle that consists of initiation, elongation, and termination stages. Coupling of the RNA processing to transcription not only coordinates timely and orderly recruitment of the RNA-processing factors but controls fidelity, efficiency and regulation of the RNA-processing reactions^2–6^. Regulation of RNA processing determines what type of RNA is made, its abundance and function. Defective RNA processing is linked to disease in humans including neurodegenerative disorders and cancer^4,7–9^. Yet, the mechanistic understanding of how Pol II transcription controls RNA processing is limited.

Shortly after transcription initiation, messenger RNA precursor (pre-mRNA) and ncRNAs undergo 5’ capping when m^7^G is attached to the 5’end of the RNA emerging from Pol II. During elongation stage, the pre-mRNA splicing occurs when non-coding introns are excised, and exons joined by the spliceosome machinery. At the 3’-end of genes, the cleavage of nascent pre-mRNA (which is polyadenylated) is linked to Pol II dislodgement and termination of transcription^10–1310,14^. A nascent transcript is cleaved and then polyadenylated by the cleavage and polyadenylation machinery which recognises polyadenylation signal (PAS) in the 3’UTR of the molecule. PAS consists of AAUAAA and GU elements recognised by the *<u>C</u>*leavage and *<u>P</u>*oly*<u>A</u>*denylation (CPA) factor (mammalian *<u>C</u>*leavage and *<u>P</u>*olyadenylation *<u>S</u>*pecificity *<u>F</u>*actor, CPSF) and *<u>C</u>*leavage *<u>F</u>*actors A and B (CFA and B in yeast or CFI and CFII in mammals)^12,13,15,16^. The RNA endonuclease subunit of the CPA (mammalian CPSF73 and yeast Ysh1) cleaves between these two elements^17^. Endonucleolytic cleavage of RNA by the CPA generates a 5’ monophosphorylated RNA end that is crucial for initiating 5ʹ–3ʹ degradation of the Pol II-associated RNA by exoribonuclease Xrn2. Xrn2 is an essential, highly conserved eukaryotic transcription termination factor that acts in a co-transcriptional manner to mediate Pol II dissociation from the DNA template to terminate transcription and facilitate Pol II recycling^18,19^. However, the key aspects of this process, including the mechanistic understanding of the steps involved and how they are coordinated in the context of transcription remain obscure. In contrast to pre-mRNA, many non-coding transcripts do not undergo splicing. They rely on the machinery distinct from the CPA, called Integrator that executes endonucleolytic cleavage of nascent RNA. In contrast to CPA, Integrator lacks poly(A) polymerase activity^20^. Following Integrator mediated cleavage, the exonucleolytic 3’ to 5’ RNA exosome complex trims the 3’end of the stable snRNAs to generate mature ncRNA or degrades transient species of ncRNA^21^.

Timely recruitment of RNA processing factors in a stage-specific manner during Pol II transcription is coordinated by the differential phosphorylation of the C-terminal domain (CTD) of Rpb1, the largest catalytic subunit of the 12 subunit Pol II complex^4,22^. Rpb1 CTD consists of multiple repeats (29 in fission yeast and 52 in mammalian cells) of the heptad consensus sequence Y_1_S_2_P_3_T_4_S_5_P_6_S_7_ that can be phosphorylated on five residues (tyrosine 1 – Y1, serine 2 – Ser2, threonine 4 – Thr4, serine 5 and 7 – Ser5 and Ser7). Phosphorylation of Ser5 by the kinase subunit of TFIIH, Cdk7, at the beginning of transcription is critical for the recruitment of the capping machinery^23–26^. Phosphorylation of Ser2 by Cdk9 kinase subunit of pTEFb coordinates recruitment of factors involved in pre-mRNA splicing and 3’end formation^27–31^. Phosphorylation of Thr4 and Tyr1 residues has also been linked to splicing and 3’end formation, although the exact role of these modifications is not fully clear^32–36^. Phosphorylation of the specific residues is reversed by the phosphatases. Fcp1 removes the phosphate group from Ser2 at the end of the genes^37–39^. Major changes in the phosphorylation status of Pol II happen when polymerase traverses through PAS. This is controlled via the action of PP1 and Ssu72 phosphatases interacting with the CPA factors^40,41^. Ssu72 de-phosphorylates Ser5 on the consensus repeats of Pol II CTD^42–44^ whereas protein phosphatase PP1 show species-specific differences towards CTD (targeting Ser5-P in mammals, Tyr1-P in budding and Thr4-P in fission yeast) and PP2A targets Ser5-P CTD^36,40,49,41–48^. However, recent studies demonstrated that mutations abolishing individual phospho-sites on CTD have no profound effect on RNA processing apart from capping that strictly depend on the recruitment of capping enzymes by Ser5-P of CTD^26,33,34,36,44,50^. Also, the linear pattern of CTD phosphorylation cannot explain the frequent occurrence of premature transcription termination caused by recruitment of CPA to cryptic PAS signals located at the 5’end of the genes despite low Ser2 levels in the beginning of transcription suggesting that there may be additional mechanisms in place that coordinate pre-mRNA-processing during transcription.

The speed of Pol II progression during the transcription cycle also contributes to pre-mRNA processing by mediating the selection of the *cis* elements by the RNA processing factors. For example, mutations in the Pol II catalytic centre that affect its processivity were demonstrated to induce defective RNA processing. Slow Pol II mutants facilitate inefficient RNA splicing events and selection of the proximal 3’end processing elements^51–55^. In contrast, fast Pol II shows intron inclusion and 3’ UTR lenghtening^52^.

Spt5 is a key essential factor that controls Pol II processivity^56–59^. With Spt4, it forms the DSIF complex (DRB, 5,6-Dichloro-1-β-D-ribofuranosylbenzimidazole-sensitivity-inducing factor). Spt5 binds to Pol II after initiation and travels with the Pol II elongation complex throughout the transcription cycle, where it is required for Pol II pausing or productive elongation^59–61^. Spt5 is one of few transcription factors functionally and structurally conserved between eukaryotes and bacteria reflecting its important role in transcription^62,63^. Eukaryotic and bacterial Spt5 share (*<u>N</u>*us*<u>G</u>*) *<u>N</u>*-terminal (NGN) and one *<u>K</u>*yprides, *<u>O</u>*uzounis, *<u>W</u>*oes (KOW) domains. In addition, Spt5 has evolved a negatively charged N-terminal region, several additional KOW domains (KOW1-5 in yeast and 1-7 in humans) and repetitive C-terminal region (Figure 1A). Although many aspects of the Spt5 functions in transcription are not fully understood, recent structural studies provided insights into Spt5 interaction with Pol II. The multidomain structure of Spt5 allows it to make numerous contacts with Pol II, nascent RNA, upstream DNA, and non-template DNA strand^64,65^. Spt5 can either support the paused state of polymerase^64,66,67^ or promote productive elongation by stabilising interaction around the DNA clamp^65,68–70^. In addition, Spt5 is also a subject to phosphorylation by Cdk9 (within CTR repeats and KOW4-5 linker)^60,71^, which plays a key role in the regulation of Pol II transcription. Unphosphorylated Spt5 is associated with the Pol II pausing observed downstream of the promoter region in eukaryotes^72–74^. Formation of the RNA clamp comprising KOW4-5 domains and recruitment of *<u>N</u>*egative *<u>E</u>*longation *<u>F</u>*actor (NELF) via unphosphorylated CTR facilitate stabilisation of the pause^75,76^. Phosphorylation of NELF and Spt5 by Cdk9 leads to the dissociation of NELF and conversion of Spt5 into a positive elongation factor resulting in the release of the promoter-proximal Pol II into productive elongation, which is the key regulatory checkpoint during Pol II transcription^60,75,77^. Currently, how phosphorylation of Spt5 contributes to Pol II processivity is not fully understood. One possibility could be that phosphorylated Spt5 repeats mediate recruitment of factors during transcription similar to what is described for Pol II CTD. Indeed, in agreement with this, recruitment of the PAF1 (the *<u>P</u>*olymerase-*<u>A</u>*ssociated *<u>F</u>*actor 1) elongation complex is facilitated by the interaction of its subunit (Rtf1) with phosphorylated Spt5. On the other hand, Spt5 CTR is de-phosphorylated at the end of the genes by PP1 phosphatase (Dis2 in fission yeast) and this is required for the efficient termination of Pol II transcription in yeast and mammalian cells^36,53,78^. Failure to de-phosphorylate Spt5 upon loss of PP1 activity or introduction of mutations in Spt5 CTR repeats mimicking constitutive phosphorylation leads to delayed Pol II dislodgement and accumulation of Pol II downstream of PAS^36,53,60,78,79^.

**Figure 1.**
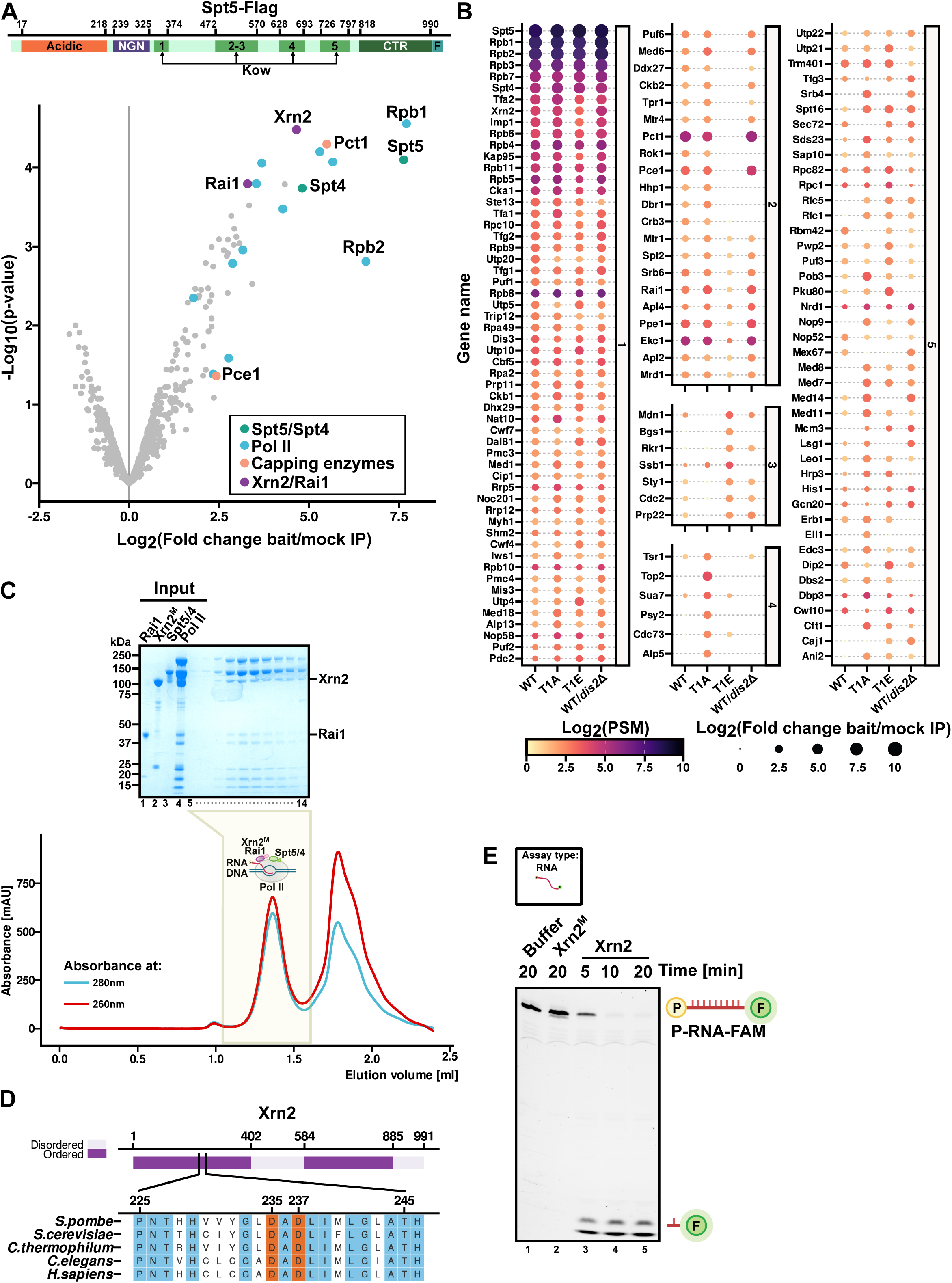
Xrn2-Rai1 complex interact with Spt5-Spt4 containing Pol II transcription complexes. (A) Volcano plot highlighting proteins enriched in Spt5-Flag purifications. The top of the figure depicts the Spt5 domain organisation. (B) Xrn2 associates with Spt5 complexes in a phosphorylation-independent manner. Proteins enriched in purifications of Spt5 from WT Spt5, T1A, T1E and cells lacking PP1 (Dis2 phosphatase) are presented as dot plot and circle size reflects a number of peptides and colour corresponds to enrichment over mock IP. Proteins are grouped according to enrichment (panel 1 – present in all, 2 – decreased in T1E or *dis2*Δ, 3 – reduced in T1A or WT, 4 – more abundant in T1A, 5 – unassigned). Metabolic enzymes and ribosomal proteins were filtered out. (C) Xrn2-Rai1-Spt5/4-Pol II-DNA/RNA form a stable complex. Scaffold 1 used is listed in Table S2 (compare Figure S2A) as previously published^65^. The reconstituted complex was subjected to size exclusion chromatography (Superose 6) and analysed by SDS-PAGE. (D) Xrn2 alignment shows conserved catalytic residues that have been mutated in Xrn2^M^ (D237A for *in vitro* or D235A for *in vivo* experiments). (E) *In vitro* Xrn2 RNA degradation assay. 3’ FAM labelled 5’-monophosphate-RNA (P-RNA-FAM) substrate was incubated with either Xrn2, Xrn2^M^ (D237A) or buffer for the indicated time. Intact RNA substrate and degradation products (indicated on the side) were resolved on 10% 8M Urea-PAGE gel.

The central assumption in the field is that after endonucleolytic cleavage of the nascent transcript by the CPA, the 5’-3’ RNA exoribonuclease Xrn2 degrades RNA and chases the transcribing Pol II to dismantle polymerase from DNA. It was recently demonstrated that de-phosphorylation of Spt5 is associated with deceleration of Pol II speed downstream of PAS^53^. This suggested that the key function of PP1-mediated Spt5 de-phosphorylation is to allow Xrn2 exonuclease to catch up with the transcribing Pol II. Interestingly, recent studies indicated that depletion of Spt5 also leads to the increased readthrough transcription and delayed termination in mammalian cells^60,79^.

Here, we employ transient transcriptome sequencing (TT-seq)^80,81^ to assess the impact on transcription upon acute degron-mediated depletion of Spt5 in fission yeast. We report widespread transcription elongation defects upon acute Spt5 loss observed for most protein-coding genes. At some protein-coding TUs loss of Spt5 leads to the pronounced signal in promoter-proximal region with concomitant decreased read density in the gene body suggesting that Pol II is either retained at the promoter or undergoes premature transcription termination in the absence of Spt5. Our analyses revealed that this cluster of TUS shows enrichment in T-tracks downstream promoters. Interestingly, our observations are consistent with recent reports demonstrating that Pol II requires help from Spt5 and the U1 complex of the spliceosome to transcribe through T-rich or A/T-rich stretches. This type of sequences might be a difficult template for Pol II inducing pausing and transcription termination^82,83^.

Additionally, we demonstrate that loss of Spt5 also leads to failed pre-mRNA splicing, 3’end processing and delayed transcription termination genome-wide. This is accompanied by a global decrease in transcription rate observed upon acute loss of Spt5 suggesting that Pol II complexes that have undergone transition into elongation from the promoter-proximal pausing in the absence of Spt5 are not competent to support pre-mRNA processing. Hence, we propose that Spt5 is involved in the quality control of Pol II complexes before entry into elongation. In contrast to protein-coding genes, transcription of multiple non-coding TUs is increased in Spt5-depleted cells suggesting that Spt5 plays a role in attenuation of non-coding transcription.

Here, we employ biochemical reconstitution, cross-linking mass spectrometry, cryo-EM, *in vitro* degradation assays and analyses of nascent transcriptome to demonstrate that formation of the stable complex between Xrn2, its interacting partner Rai1 (DXO in human), DSIF (Spt5-Spt4, Spt5/4) and 12 subunit Pol II DNA-RNA scaffold is a prerequisite for optimal exoribonucleolytic activity of Xrn2. In this context Spt5 is important for full activation of Xrn2 facilitating efficient transcription termination. We demonstrate that Xrn2 and Rai1 are highly enriched in Spt5 native purifications together with well-documented Spt5 interacting proteins such as Pol II subunits and capping enzymes^60,84^. These insights bring a new perspective on the accepted view that Xrn2 chases down still transcribing Pol II after cleavage, and rather suggest that Xrn2 functions as a part of Pol II complex in a collaboration with Spt5. In agreement with our findings, recent studies that employed transcriptome analyses mapping Xrn2 entry sites and PAS usage proposed that Xrn2 acts in close association with Pol II throughout the transcription cycle and engages with the 5’end of the nascent RNA immediately after it is cleaved by CPA^52,85^. We show that Spt5 activates Xrn2 independently of its phosphorylation by Cdk9. Based on our data, we conclude that Spt5 plays a dual role during transcription termination where it coordinates kinetics of Pol II transcription and RNA degradation to ensure timely termination of transcription and accurate gene expression. Both Spt5 and Xrn2 stand out in that they are highly conserved and function in the regulation of other polymerases such as Pol I which implies that the link between these proteins may also be universally conserved^70–74^. Interestingly, NusG, the bacterial homologue of Spt5, is required for the recruitment and activity of Rho helicase, the factor that terminates transcription in bacteria further emphasising the conceptual parallel between eukaryotic and bacterial mechanisms involved in termination of transcription^86^.

## RESULTS

### Xrn2 forms a pre-termination complex with DSIF-Pol II

Spt5 is phosphorylated by Cdk9 on threonine 1 (Thr1, T1) within its C-terminal region (CTR) composed of the repeats of a nonapeptide motif of the consensus sequence T^1^PAWNSGSK. Previous studies suggested that phosphorylation of Spt5 facilitates Pol II escape from the promoter-proximal pausing and supports productive elongation, whereas removal of phosphorylation from Spt5 by PP1 phosphatase contributes to transcription termination in yeast and mammalian cells^36,53,60,72,78,87,88^. To gain insight into how phosphorylation mediates Spt5 function in transcription, we undertook a proteomic approach. We purified a tagged Spt5 from the wild-type (WT) fission yeast strain and a strain lacking one of the PP1 variants - Dis2 (D*dis2*). In parallel, we also purified mutated Spt5 where T1 in all repeats was either replaced by alanine (T1A) or glutamate as a phosphomimic (T1E). Surprisingly, analyses of the Spt5 purifications by mass spectrometry indicated a high enrichment of the components of the transcription termination machinery: the 5’-3’ exonuclease Xrn2 and its interacting partner Rai1 in addition to known interactors of Spt5 such as Spt4, components of the RNA 5’-capping machinery (RNA-triphosphatase, Pct1 and guanylyl-transferase, Pce1) and subunits of the Pol II (Figure 1A, B - panel 1, Table S1). However, the high enrichment of Xrn2 in all Spt5 purifications suggests that this factor interacts with Spt5-associated complexes in a phosphorylation-independent manner (Figure 1B). In contrast, components of the capping machinery, Pct1 and Pce1, were not detected in the purification of Spt5 T1E mutant (Figure 1B-panel 2) while enriched in the purifications from other mutants and WT Spt5. These data are consistent with previous studies demonstrating that these enzymes specifically recognize unphosphorylated Spt5 and their binding to Spt5 is antagonised by Thr1 phosphorylation^89–91^. At the same time, Rtf1 subunit that interacts with phospho-Spt5^92–95^ as well as other components of the PAF1 complex were not present in either WT nor T1E Spt5 purifications suggesting that the comparative proteomics approach can be utilised to identify stable interactions but may not be applicable to identify more transient interactors (Figure 1B, Table S1).

Next, we employed a fully defined *in vitro* system to test whether Xrn2/Rai1 heterodimer interacts directly with Spt5 and Pol II. To this end, we purified native Pol II via 3xFlag-tagged Rpb9 using affinity chromatography followed by ion exchange (Figure 1C, SDS-PAGE lane 4). The Spt5/4 (DSIF) complex was co-expressed in bacteria (Figure 1C, SDS-PAGE lane 3). The functionality of the reconstituted DSIF in stimulating transcription elongation was validated using a promoter-independent transcription elongation assay^96,97^. To achieve that, native Pol II was assembled on a dsDNA/RNA transcription bubble in the presence or absence of Spt5/4, and transcription was started by the addition of ribonucleotides (Figure S1A, B and C, Table S2). Analysis of the products of the transcription reaction demonstrated that Spt5/4 stimulated Pol II processivity and reduced the number of pausing events (Figure S1C, compare lanes 2 and 5) in agreement with the previously published work^98,99^.

To study the nucleolytic activity of Xrn2 we expressed WT and catalytic mutant (Xrn2^M^, D237A), in which a conserved aspartic acid located in the catalytic pocket of the enzyme was replaced with alanine (Figure 1D)^18,42,54,85,100,101^. To obtain a high yield of stable protein, both constructs were expressed without the C-terminal part (1-885 aa of Xrn2) as previously reported^101^. The ability of the WT and mutated enzyme to degrade RNA was studied using 3’ FAM-labelled RNA with 5’ monophosphate (Figure S1D, P-RNA-FAM, Table S2). This RNA mimics the Xrn2 substrate during the transcription termination process, which corresponds to the product of endonucleolytic cleavage of the pre-mRNA by the cleavage/polyadenylation machinery. Incubation of C-terminally truncated WT Xrn2 but not Xrn2^M^ with the RNA substrate resulted in gradual disappearance of the full-length RNA and accumulation of short degradation products after 5 min of incubation (Figure 1E, compare lane 1 to lanes 3-5). Having established that Xrn2^M^ does not possess catalytic activity, we aimed to reconstitute a complex between Xrn2, Rai1, DSIF and Pol II with a DNA-RNA scaffold^65^ representing *an in vitro* transcription elongation bubble. We used Xrn2^M^ to avoid degradation of the Pol II bound RNA and destabilisation of the complex. Reconstituted Pol II-DSIF-Xrn2/Rai1 complex was then subjected to size exclusion chromatography demonstrating that Xrn2/Rai1 factors stably associate with elongating Pol II/DSIF. We refer to this complex as a Pre-Termination Complex (PTC) (Figure 1C, SDS-PAGE lanes 7-14).

### Xrn2 directly interacts with Spt5 and Rpb2-Rpc10 interface of Pol II

To further investigate if the interaction between Spt5/4 and termination factors depends on Pol II, we performed pull-down assays. Rai1 was immobilised and incubated either with Xrn2 alone or Xrn2+Spt5/4 (Figure 2A). This experiment revealed that Spt5/4 can bind Xrn2/Rai1 independently of Pol II. To test the Spt5/4 requirement for the interaction of Xrn2 with Pol II, we prepared Pol II complexes immobilised on anti-Flag beads via Rpb9 with or without nucleic acids and ±Spt5/4 (Figure S2A). This experiment revealed that Xrn2 can also interact with Pol II assembled on the DNA-RNA scaffold independently of Spt5/4 and Rai1 (Figure 2B, lanes 9-10, 12-13). However, in the absence of nucleic acids, the association of Xrn2 with the Pol II required the presence of Rai1 (Figure 2B, compare lanes 4 to 3 and 6 to 7).

**Figure 2.**
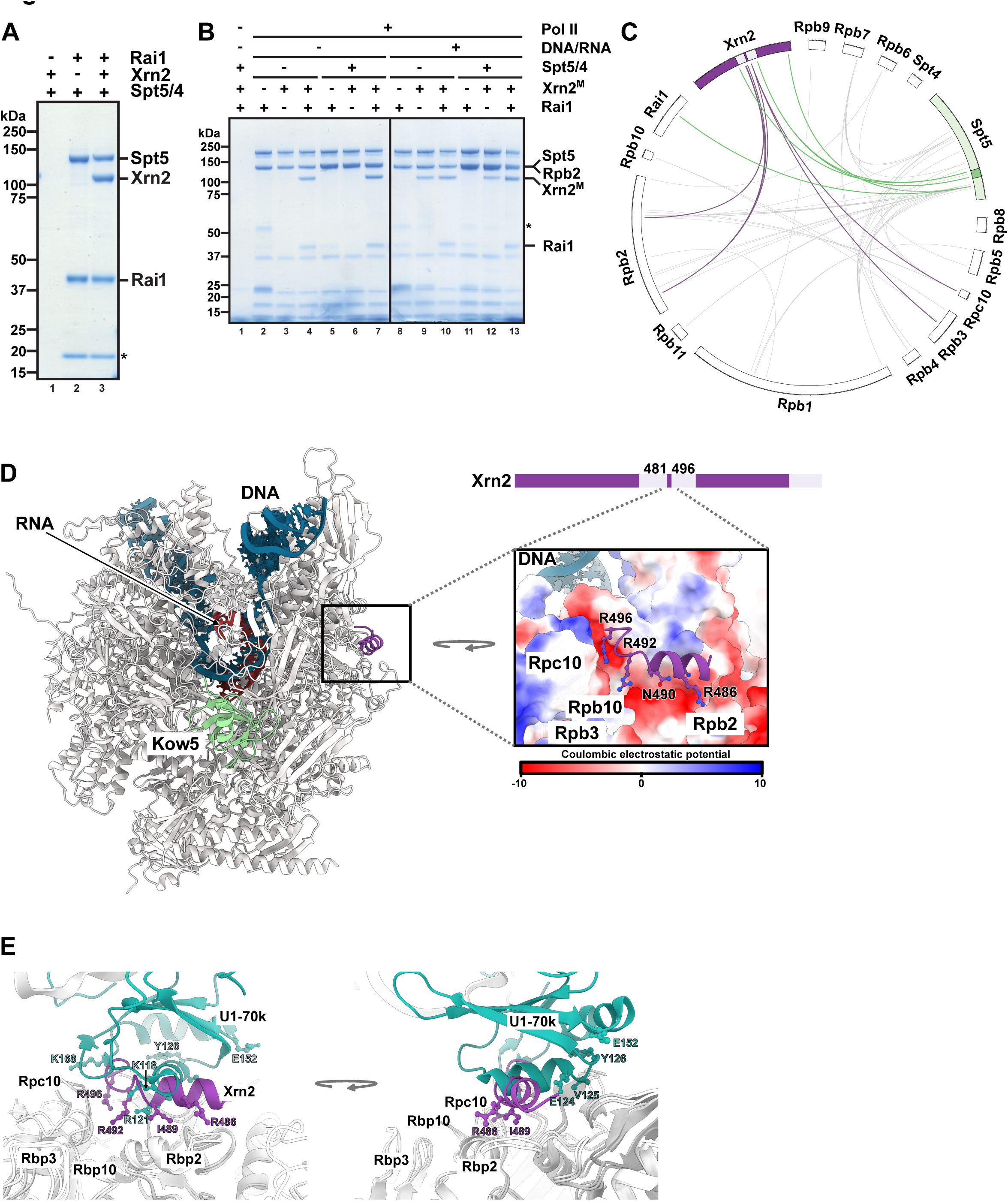
Xrn2 interacts with Pol II and contacts Spt5 within KOW5 region. (A) Spt5 interacts with Xrn2/Rai1. Rai1 was immobilised on beads and challenged with Xrn2 or Xrn2/Spt5/4. The asterisk indicates an unspecific degradation product. (B) Xrn2 can interact with Pol II independently of Spt5. Pol II complex with or without nucleic acids was immobilised on beads (Figure S2A, Table S2 - scaffold 1). Complexes were incubated with indicated proteins and washed and mixtures were resolved on SDS-PAGE. The asterisk marks antibody heavy chain. (C) Xrn2-Rai1-Spt5/4-Pol II-DNA-RNA scaffold complex cross-linking with BS3 coupled to mass spectrometry. Inter-protein crosslinks are shown as lines. Crosslinks between Xrn2 and Pol II are coloured purple and Spt5-Xrn2/Rai1 in green. (D) Cryo-EM model of the Xrn2-Rai1-Spt5/4-Pol II-DNA-RNA scaffold complex. RNA is shown in red, DNA in blue, the Spt5 KOW5 domain in green and the α-helical loop of Xrn2 in purple, which docks enzyme to the negatively charged surface of Pol II (zoomed area). (E) Xrn2 is anchored to the same Pol II region as U1-70k from splicing machinery. Pol II-U1 snRNP complex (PDB: 7B0Y)^104^ was superimposed on the Pol II-Spt5-Xrn2 structure.

To further identify interacting interfaces within the PTC complex, we performed chemical crosslinking of the reconstituted Pol II-Spt5/4-Xrn2^M^-Rai1-DNA-RNA complex with bis (sulfosuccinimidyl)suberate (BS3) (Figure S2A and S2B, lane 8) and analysed cross-linked peptides using mass spectrometry (Table S3). A cluster of crosslinks is observed in the middle region of Xrn2 contacting Pol II (Rpb2, Rpb3 and Rpc10) as well as KOW5 and CTR of Spt5 (Figure 2C). The region of Xrn2 engaged in interactions with Pol II is located between two parts of the enzyme that constitute its single catalytic domain. This region is not visible in the published high-resolution structure of Xrn2 with Rai1 and is predicted to be unstructured^101^. To gain a deeper understanding of the interactions within the reconstituted PTC complex, we subjected Pol II-DSIF-Xrn2^M^/Rai1/DNA-RNA complex to glutaraldehyde crosslinking (Figure S2A and S2B, lane 9) and cryo-electron microscopy (cryo-EM). We were able to obtain high-resolution 3D-reconstitutions of Pol II (at 2.67Å, Table S4) with nucleic acid scaffold and additional densities next to the RNA exit channel (corresponding to KOW5 of Spt5) and the region in the vicinity of Rpb2/Rpb10/Rpc10 (Figure S2C). In these reconstructions density corresponding to Rpb4/7 is missing (Figure S2C). Crosslinking data and AlphaFold^102,103^ guided the modelling of the additional density, which corresponds to Xrn2 residues 481-496 (Figure 2D, S2C and S2D). We suggest that Xrn2 uses a small alpha-helical region embedded in the unstructured segment to anchor the enzyme to Pol II. These interactions are mostly ionic with arginines (R492 and R496) of Xrn2 docking to a negatively charged region within Pol II (Figure 2D, E). Additionally, isoleucine 489, located in the middle part of the Xrn2 peptide contacts a hydrophobic/negatively charged stretch of Rpb2 (Figure 2D and E). Although we observe only a small helical region from Xrn2 bound to Pol II, given the crosslinking data and the flexible nature of this region, the interaction surface may be larger and most likely transient. Despite the low sequence conservation of the unstructured Xrn2 region, AlphaFold predicts that human and mouse Xrn2 enzymes contain small helical segments similar to the fission yeast counterpart with arginine residues present (Figure S2E). These data suggest that Xrn2-Pol II would form similar interfaces in higher eukaryotes and the interaction might be driven by ionic forces. Interestingly, the U1 complex of the splicing machinery binds to the same region on the Pol II (Figure 2E)^104^. U1-70K contacts Pol II using positively charged lysines and arginines (i.e. R121 and K118) but the orientation of the helix is rotated by 90° relative to the Pol II-interacting helix of Xrn2 (Figure 2E). Therefore, splicing and termination machinery may compete for binding to the same Pol II surface.

### Spt5 stimulates exonucleolytic activity of Xrn2

To further understand whether there is a functional link between Spt5/4 and Xrn2, we tested its effect on the 5’ to 3’ exoribonucleolytic activity of C-terminally truncated Xrn2 on 5’ monophosphorylated substrate with the FAM label at the 3’ end (Figure S1D, P-RNA-FAM). Analyses of the reaction products by UREA-PAGE revealed more efficient degradation of the substrate RNA by Xrn2 in the presence of Rai1 in agreement with previous study proposing stabilization of Xrn2 by Rai1^101^ (Figure S3A, compare lanes 3-5 and lanes 12-14). Strikingly, the addition of Spt5/4 had a stimulatory effect on RNA degradation by Xrn2 and appeared to override the dependency of Xrn2 on Rai1 (Figure S3A, compare lanes 3-5 and lanes 15-20). To better approximate the natural substrate of Xrn2 in the context of Pol II termination, we also assessed the effect of Spt5/4 on Xrn2 activity by performing RNA degradation assays on P-RNA-FAM that we assembled as a part of DNA-RNA scaffold with Pol II (Figure S3B). Consistent with our previous observations, Spt5/4 also stimulated Xrn2 activity on the RNA substrate associated with Pol II (Figure 3A, compare lanes 5 and 7 to 6 and 8). We conclude that Spt5 can stimulate Xrn2 exoribonucleolytic activity towards RNA and suggest that this is relevant during transcription.

**Figure 3.**
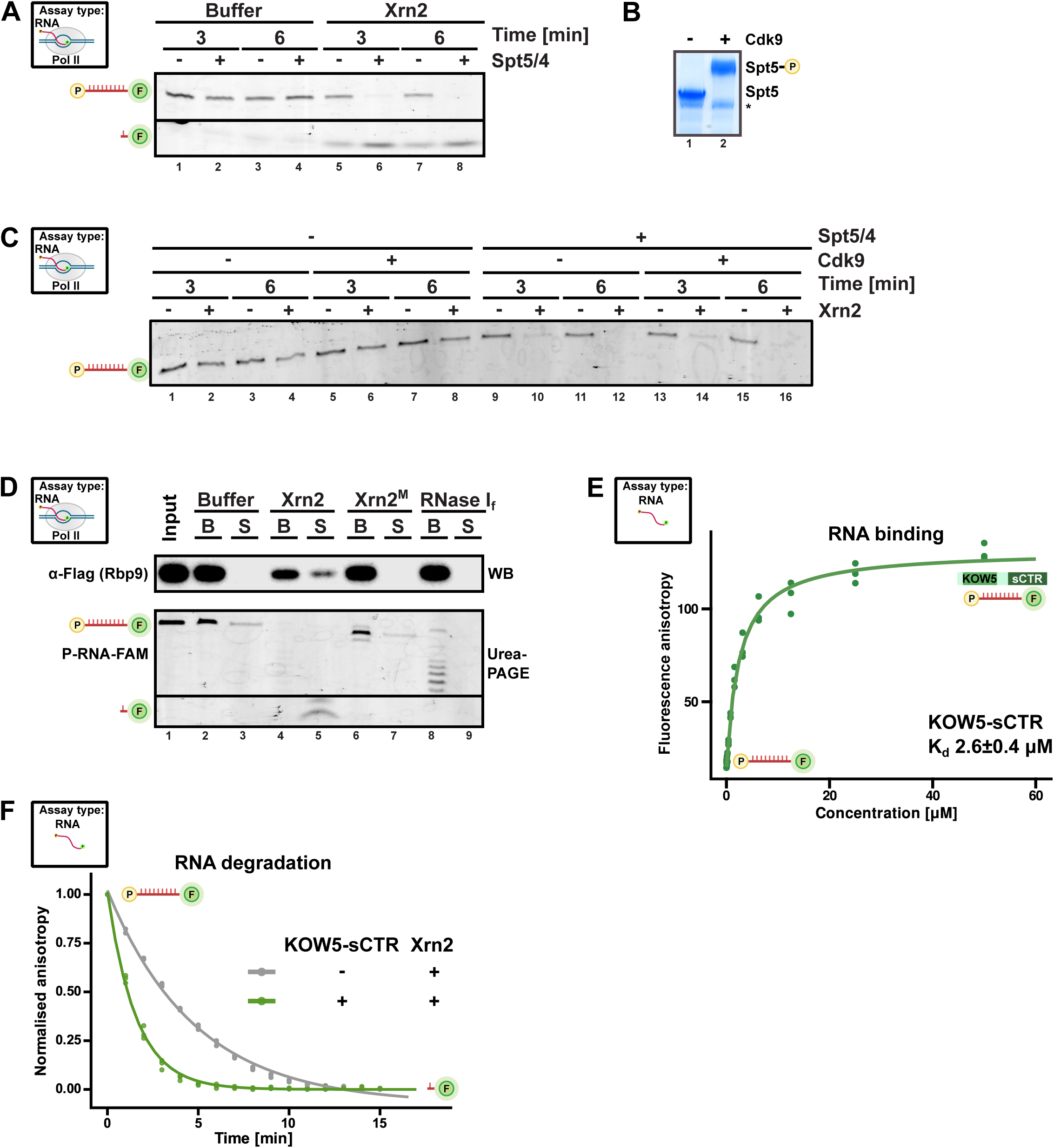
Spt5 stimulates Xrn2-dependent RNA degradation. (A) Spt5 can stimulate Xrn2 exoribonucleolytic activity towards 3’FAM labelled RNA in the context of the Pol II complex. Complexes were immobilised on beads using biotinylated non-template DNA (compare Figure S3B). (B) Spt5 phosphorylation by the recombinant Cdk9/Cyclin T1 complex resolved by the phos-tag gel. (C) Spt5 stimulates RNA exoribonucleolytic activity of Xrn2 independently of phosphorylation (compare Figure S3B). (D) Degradation of nascent RNA by Xrn2 but not RNase I_f_ dislodged Pol II. Transcription termination was performed using an *in vitro* system with Pol II immobilised on beads via biotinylated non-templated DNA (compare Figure S3C). S - corresponds to the supernatant fraction, B is beads fraction. (E) The affinity of KOW5-sCTR was evaluated using a polarisation anisotropy assay. 3’FAM labelled 5’-monophosphate-RNA substrate was titrated with increasing amounts of purified protein. (F) KOW5-sCTR (Spt5 region 720 to 874 aa, N-terminal His-tag) stimulates degradation by Xrn2. Fluorescence polarisation anisotropy assay comparing RNA degradation kinetics of Xrn2 alone or KOW5-sCTR (compare Figure S3E, F).

We next aimed to determine whether phosphorylation contributes to Spt5 ability to stimulate RNA degradation by Xrn2. To phosphorylate Spt5, we reconstituted the Cdk9/Pch1 (cyclin T1) complex and performed an *in vitro* phosphorylation reaction by incubating kinase with Spt5/4. Efficient phosphorylation of Spt5 was confirmed by the shift in the Spt5 mobility using a phos-tag gel (Figure 3B, compare lanes 1 and 2). Next, the activity of Xrn2 towards RNA substrate in the context of Pol II was assessed in the presence of non-phosphorylated or phosphorylated Spt5/4 (Figure 3C). Phosphorylated and non-phosphorylated Spt5 stimulate Xrn2 activity to a similar extent (Figure 3C, compare lanes 9-12 to lanes 13-16) implying that Spt5 acts independently of phosphorylation status in stimulating Xrn2 which is consistent with our Spt5 proteomic data (Figure 1B).

### Exoribonucleolytic activity of Xrn2 is important for dislodgement of Pol II from DNA and transcription termination

Previous studies performed in *Saccharomyces cerevisiae* (*S. cerevisiae*), and human demonstrated that mutation of the active site of Xrn2 (D235) leads to a global shift of Pol II occupancy further downstream of the PAS suggesting that exoribonucleolytic activity of Xrn2 is required for transcription termination *in vivo*^54,100^. To test whether the ability of Xrn2 to degrade RNA is required for the dislodgement of Pol II from DNA we set up an *in vitro* transcription termination assay (Figure S3C)^96,105^. Here, Pol II was loaded onto a DNA-RNA scaffold and immobilised on streptavidin beads followed by incubation with C-terminally truncated Xrn2 WT or Xrn2^M^. The Pol II dislodgement from the DNA template was monitored by assessing Rpb9 levels in the supernatant upon incubation with Xrn2. Only WT Xrn2 led to the Pol II presence in the supernatant which correlated with the accumulation of RNA degradation products in supernatant and the disappearance of RNA from the beads (Figure 3D, lanes 4 and 5). In contrast, no Pol II release or RNA degradation was observed upon incubation with mutant Xrn2^M^ (Figure 3D, lanes 6, 7). As a control, we also incubated Pol II with RNase I_f_ which has endonucleolytic activity towards RNA. Incubation with RNase I_f_ did not result in Pol II release and led to the accumulation of RNA fragments corresponding to the size of the RNA protected by Pol II (16-18nt) (Figure 3D, lanes 8, 9). Taken together, we confirm that 5’ to 3’ exoribonucleolytic activity of Xrn2 directly contributes to Pol II dislodgement from the DNA template.

### Mapping the region of Spt5 that contributes to interaction with Xrn2 and its activation

The observation that Xrn2 crosslinks to Spt5 in the vicinity of the KOW5 domain (Figure 2C) prompted us to examine whether this region is sufficient for the stimulation of Xrn2 enzymatic activity. We expressed and purified KOW5 with part of the CTR region (KOW5-sCTR) that includes amino acids 720 to 874 of Spt5 (Figure S3D). To test whether the addition of KOW5-sCTR affects Xrn2 RNA degradation kinetics we performed reactions in the presence and absence of this domain. To quantitatively study RNA degradation kinetics we employed an *in vitro* assay based on fluorescence anisotropy (FA) measurements (Figure S3E). Xrn2 degradation of the substrate P-RNA-FAM releases the fluorescent dye (FAM), and it can be monitored as a decrease of the anisotropy over time. Since Spt5 can interact with DNA and RNA we evaluated the binding of KOW5-sCTR to RNA using a fluorescence polarisation assay, which indicated that this construct has a modest affinity towards RNA (Figure 3E). The presence of KOW5-sCTR stimulated Xrn2 activity as the half-life of normalised anisotropy (a proxy for substrate half-life) decreased ∼3-fold in the presence of the Spt5 construct (Figure 3F). The stimulatory effect of KOW5-sCTR was comparable to the full-length Spt5 (Figure S3F). The contribution of Spt5 and KOW5-sCTR binding to RNA to the anisotropy was considered when assessing their effect on the RNA degradation kinetics of Xrn2. We conclude that Spt5 stimulates Xrn2 exoribonucleolytic activity via the region containing the KOW5 domain.

### Loss of the exoribonucleolytic activity of Xrn2 leads to profound global dysregulation of transcription

Next, we examined the contribution of Xrn2 catalytic activity to transcription. To generate a strain expressing Xrn2^M^, we introduced a genomic copy of V5 tagged *xrn2* gene with the D235A mutation in a strain expressing plant-derived protein Tir1 (Transport Inhibitor Response 1, the auxin receptor F-box protein) and WT miniAIDx3-Flag-tagged Xrn2 protein^106^. In parallel, we also generated a control strain that constitutively expressed WT V5-tagged Xrn2 to complement for auxin-depleted AID-tagged Xrn2 protein. We verified that V5-tagged WT and mutant Xrn2 proteins were expressed (Figure 4A, lanes 3-6) and localized to the cell nucleus (Figure S4A). AID-tagged Xrn2 protein was efficiently depleted after growing cells for 2h in the presence of auxin (Figure 4A, compare lanes 1 and 2, 3 and 4, 5 and 6). Xrn2 and its catalytic activity essential for cell viability (Figure S4B). Expression of WT but not the mutated version of Xrn2 fully rescued the growth of Xrn2-AID in the presence of auxin (Figure S4B). To facilitate the design of Xrn2 mutants in the future, we took advantage of this system to map the location of the nuclear localisation signal (NLS) of Xrn2 by generating a series of deletions (Figure S4C). Interestingly, deletion of the 444-575 aa Xrn2 region, although preserving the previously predicted NLS^107^ (396 to 418 aa of Xrn2), interfered with protein localisation to the nucleus. Further analyses of the Xrn2 sequence identified an additional low-scoring, putative NLS^108^ within the deleted fragment, located between 566-574 aa of Xrn2. Indeed, a mutant that only lacks the region from 444-555 aa of Xrn2 shows nuclear enrichment (Figure S4C) confirming that the Xrn2 region between 555-574 aa contains an important NLS. To initially assess the contribution of the catalytic activity of Xrn2 to termination efficiency in the cell in a quantitative manner, we harnessed the system that controls the expression of the phosphate-acquisition genes including Pho1 phosphatase^109–112^. Expression of the *pho*1 gene is controlled by premature transcription termination of lncRNA termed *prt* (*<u>P</u>*ho1-*<u>R</u>*epressing-*<u>T</u>*ranscript) (Figure S4D). The transcription termination efficiency of *prt* can be assessed by quantifying the enzymatic activity of Pho1 in a colorimetric assay which relies on the dephosphorylation of *p*-nitrophenylphosphate to *p*-nitrophenol. As predicted, acute depletion of Xrn2 leads to a more than 3-fold reduction in Pho1 activity in agreement with compromised transcription termination of *prt* lncRNA leading to repression of *pho*1 transcription (Figure 4B). Expression of WT Xrn2 can fully compensate for the loss of degron-tagged protein. In contrast, expression of the catalytic mutant following depletion of AID-tagged WT Xrn2 leads to an additional reduction in Pho1 activity compared to auxin depletion alone, which might be due to the dominant negative effect imposed by expression of Xrn2^M^. These data confirm the importance of the catalytic activity of Xrn2 for its function in transcription termination. In agreement with our data, Xrn2 exonucleolytic activity has been recently demonstrated to play a role in the control of a widespread premature transcription termination in human cells^85^ in addition to its function in Pol II dislodgement at the end of the genes.

**Figure 4.**
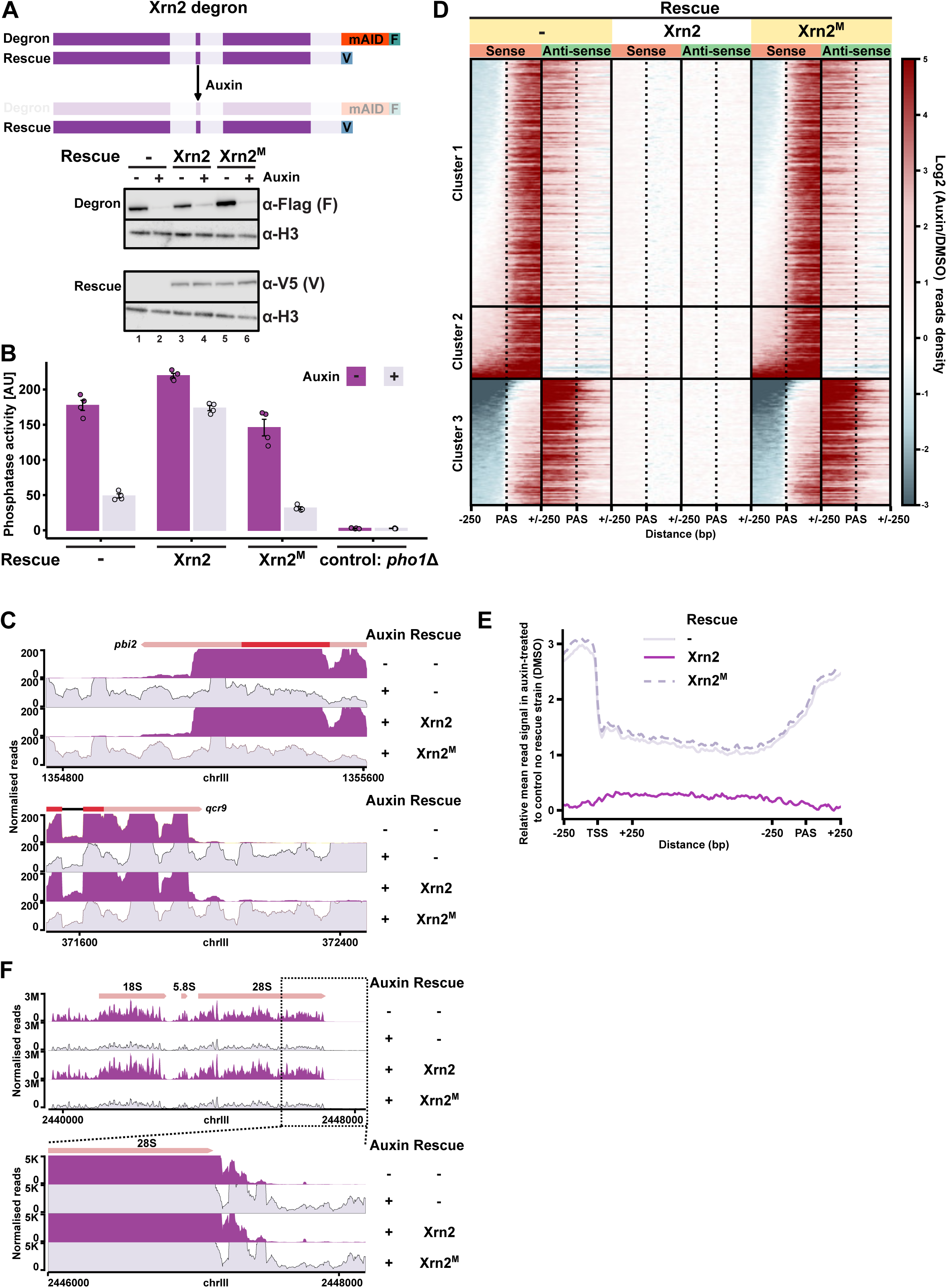
The catalytic activity of Xrn2 is indispensable for its function. (A) Auxin**-** mediated depletion of Xrn2. Additional strains were created where WT or catalytic inactive Xrn2 mutant (Xrn2^M^ - D235A) were introduced to complement depleted protein. Western blot analyses of AID-Flag tagged Xrn2 and constitutively expressing V5-tagged WT and Xrn2^M^. Histone (H3) serves as a loading control. (B) Assessment of transcription termination using endogenous phosphatase activity-based reporter system (Figure S4D). Phosphatase activity of secreted acid phosphatase Pho1 serves as a proxy for termination efficiency. Xrn2^M^ cannot complement the depletion of WT enzyme. (C) Functional analyses of transcription upon acute depletion of Xrn2 or expression of catalytically inactive Xrn2 by transient transcriptome analysis (TT-seq). Representative snapshots demonstrating severe readthrough transcription in the absence of Xrn2 or expression of catalytically inactive Xrn2. (D) Heatmap showing gene clusters differentially affected upon Xrn2 depletion or inactivation. Transcription on sense and anti-sense strands is presented as a log_2_ fold ratio to control strain (DMSO treated strain without Xrn2 complementation). Cluster 1 and cluster 2 indicate that transcription downregulation correlates with readthrough on the opposite strand. Cluster 2 includes genes that appear as upregulated. (E) Evaluation of genes in cluster 2 (Figure 4D) presented as relative metagene. Accumulation of signal before TSS indicates that upregulation is caused by global readthrough. (F) Loss of Xrn2 activity results in transcription termination defects at Pol I transcribed rRNA genes.

Having established that Xrn2 ability to degrade RNA contributed to Pol II dislodgement from DNA and mediates efficient termination of transcription of *prt* gene in fission yeast, we next set out to investigate its global impact on transcription. We performed analysis of newly synthesized RNA by spike-in normalised transient transcriptome sequencing (TT-seq)^80,81^ in strains expressing either a single copy of AID-tagged Xrn2 or a second copy of the gene (WT or Xrn2^M^) that complements auxin-depleted Xrn2-AID. To assess the successful incorporation of the nucleotide base analogue 4-thiouracil (4-tU) and subsequent biotinylation of the labelled RNA, we analysed RNA purified on Streptavidin resin from cells grown in the presence and absence of 4-tU. RNA was only enriched after pull-down when cells had been labelled with the nucleotide base analogue (Figure S4E). Having confirmed specific pull-down of newly synthetised RNA, we prepared and sequenced TT-seq libraries. Analyses of the sequencing data revealed that Xrn2 depletion leads to an increase in the reads downstream of the annotated PAS, which is expected when either mRNA 3′ end processing, transcription termination or both processes are impaired. Coding TUs that do not have another gene on the same strand within 250 bp upstream of TSS or downstream of PAS were selected for analysis (3190 out of 6952 TUs annotated^113,114^). Expression of the catalytically inactive Xrn2^M^ led to profound changes in gene expression similar to the impact on the transcriptional landscape observed when the loss of WT AID-tagged Xrn2 was not compensated (Figure 4C). This includes the dramatic increase in the reads downstream of the PAS region demonstrating a global defect in transcription termination in this mutant (Figure 4C, D, S4F).

These observations are consistent with the published ChIP-seq data that demonstrated Pol II accumulation downstream of PAS observed in Xrn2 D235A catalytic mutant and recent analyses of nascent transcription (that employed bromouridine for metabolic labelling of RNA, Bru-seq) when Xrn2 D235A expression was induced in addition to the endogenous WT Xrn2^54,85,100^. Global accumulation of the 3’-reads observed in Xrn2^M^ is accompanied by a decrease in TT-seq signal over the gene-body region for many genes (Figure 4D, clusters 1 and 3). This could be caused by the failure in transcription termination preventing the recycling of Pol II. Alternatively, readthrough transcription can interfere with the transcription of the neighbouring genes leading to their repression. This is expected when TUs are in convergent or tandem orientation. In this scenario, a decrease of signal in the gene body is expected to correlate with the increased transcription in the anti-sense orientation (Figure 4D, clusters 1 and 3 and Figure S4G). In some cases, failed termination of transcription of the upstream gene can lead to transcription interference preventing transcription initiation at the promoter of the downstream gene (Figure S4H). In contrast, some genes show increased signal in the 3’ flanking regions and across the gene body (Figure 4D, cluster 2). The upregulation of gene body signal could be due to failure to dislodge Pol II prematurely or due to a high level of readthrough transcription originating from the upstream region over the lowly expressed gene. To distinguish between these scenarios, we plotted a relative meta profile for genes in cluster 2 (Figure 4D), which indicates that this increase can be attributed to readthrough as an elevated signal upstream of the TSS is present (Figure 4E). Interestingly, acute depletion of Xrn2 as well as loss of its activity leads to an increase in readthrough transcription downstream of the genes encoding for ribosomal RNA (rRNA) suggesting that Xrn2 plays a role in the termination of Pol I transcription (Figure 4F). This is consistent with previous work demonstrating that *S. cerevisiae* Xrn2 is required for termination of transcription at Pol I transcribed rDNA arrays and implies that Xrn2 plays a universal conserved role in termination of transcription^115,116^. Together, based on combined analyses we conclude that catalytic activity is key to Xrn2 function in transcription.

### Spt5 is required for restricting non-coding transcription and ‘licencing’ of Pol II complexes at genic promoters

To study the direct contribution of Spt5 to transcription, we constructed an *Schizosaccharomyces pombe* (*S. pombe*) strain that allows for rapid, auxin-inducible degradation of Spt5 to minimise the possibility of pleiotropic secondary effects that might occur upon prolonged loss of this key essential factor. Tagging Spt5 with triple miniAID tag^106^ achieves nearly complete depletion of Spt5 within 2h (Figure 5A). To assess the contribution of Spt5 to transcription we performed spike-in normalized TT-seq in the cells treated with auxin or DMSO which is highly reproducible (Figure S5A). Upon Spt5 depletion, there is a global reduction in the RNA synthesis rate (Figure 5B, S5B) in agreement with the well-documented role of Spt5 as a key positive transcription elongation factor^60,117,118^. Interestingly, a subset of genes exhibits non-uniform distribution of the nascent RNA signal along the gene after Spt5 depletion. This class of genes is characterised by a relatively high TSS/promoter-proximal signal relative to the low read density in the gene body (n=465 coding TUs) (Figure 5C, D) and might be subject to premature termination. Further, Spt5 depletion results in a high promoter-proximal signal for some TUs in this group (Figure 5C, top panel). This is consistent with the proposed role for Spt5 in the control of the Pol II promoter-proximal pausing^117,119^ by facilitating the conversion of the elongation complex into a pause-resistant state^79^. These genes (n=465) show elevated T content with a tendency to form runs of poly-T (poly-U RNA) (Figure S5C). Transcription of poly-U stretches is known to terminate Pol III^96,120–122^ and this type of sequence bias might show a strong dependence of Pol II on Spt5 for processive elongation. Recently, budding yeast Pol II has been shown to terminate on T-tracks in the absence of Spt5 which counteracts sequence-dependent premature transcription termination^82^. Gene ontology analysis revealed that genes with a pronounced pile-up of reads at the 5’-end are enriched in categories related to stress, external stimulus responses, and regulation of Pol II in these conditions (Figure S5D, Table S5) which might indicate that genes in this category have a higher transcriptional plasticity.

**Figure 5.**
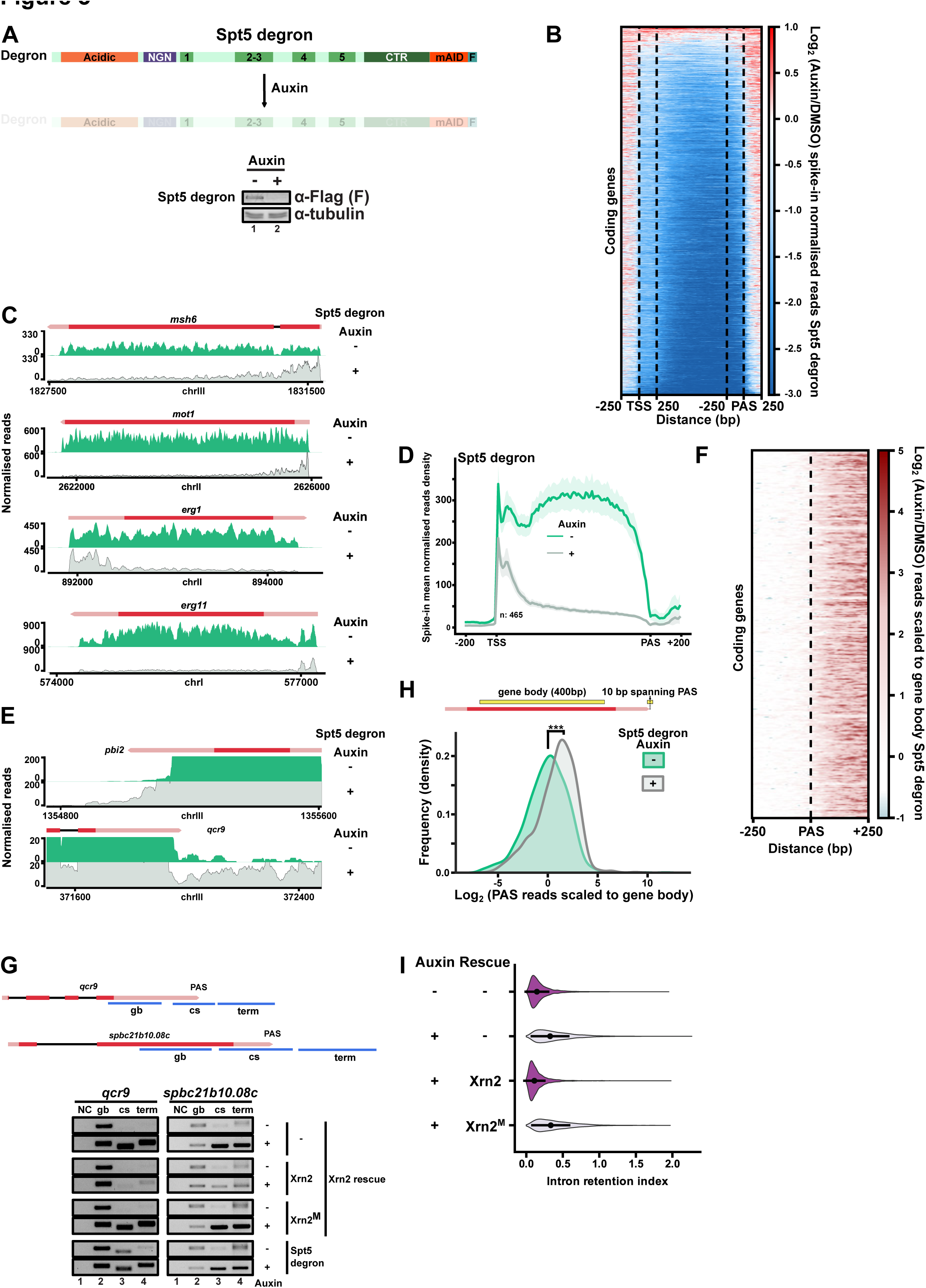
Spt5 depletion affects multiple aspects of transcription including readthrough. (A) Auxin**-**mediated depletion of Spt5. Western blot analyses of AID-Flag tagged Spt5. Tubulin serves as loading control (B) Heatmap of log_2_ ratio TT-signal between auxin and DMSO treated Spt5 degron strain. Gene body signal is scaled in the region between 250 bp after TSS and before PAS. (C) Example TT-seq genome tracks snapshot to illustrate 5’end premature transcription termination/attenuation upon Spt5 depletion. (D) Metagene comparing profiles of preselected coding genes showing premature termination/attenuation before and after Spt5 loss (shaded areas indicate 95% confidence intervals). (E) Genome tracks highlighting transcription readthrough after Spt5 depletion. (F) Genome-wide transcriptional readthrough on coding genes. The region around PAS is presented as a heatmap with a log_2_ fold ratio of auxin and DMSO treated Spt5 degron. Due to the global downregulation of transcription upon Spt5 loss, the read density was normalised to gene body (compared to Figure S4F). (G) RT-PCR analysis of readthrough and cleavage in indicated strains. Gene body – gb, cleavage site region – cs and termination zone – term are depicted on the top panel. NC refers to the negative control (PCR for gb, without reverse transcription) (H) Spt5 might contribute to 3’end processing. Experimental PAS sites^140^ were filtered to keep the most pronounced and the closest to gene end with PAS motif in 50 bp upstream sequence. Read density was calculated in a 10bp window around PAS and normalised to the gene body density. The plot shows changes in distributions before and after Spt5 depletion. (I) Violin plots showing intron retention index in Xrn2-degron strains before and after auxin treatment.

**Figure 6.**
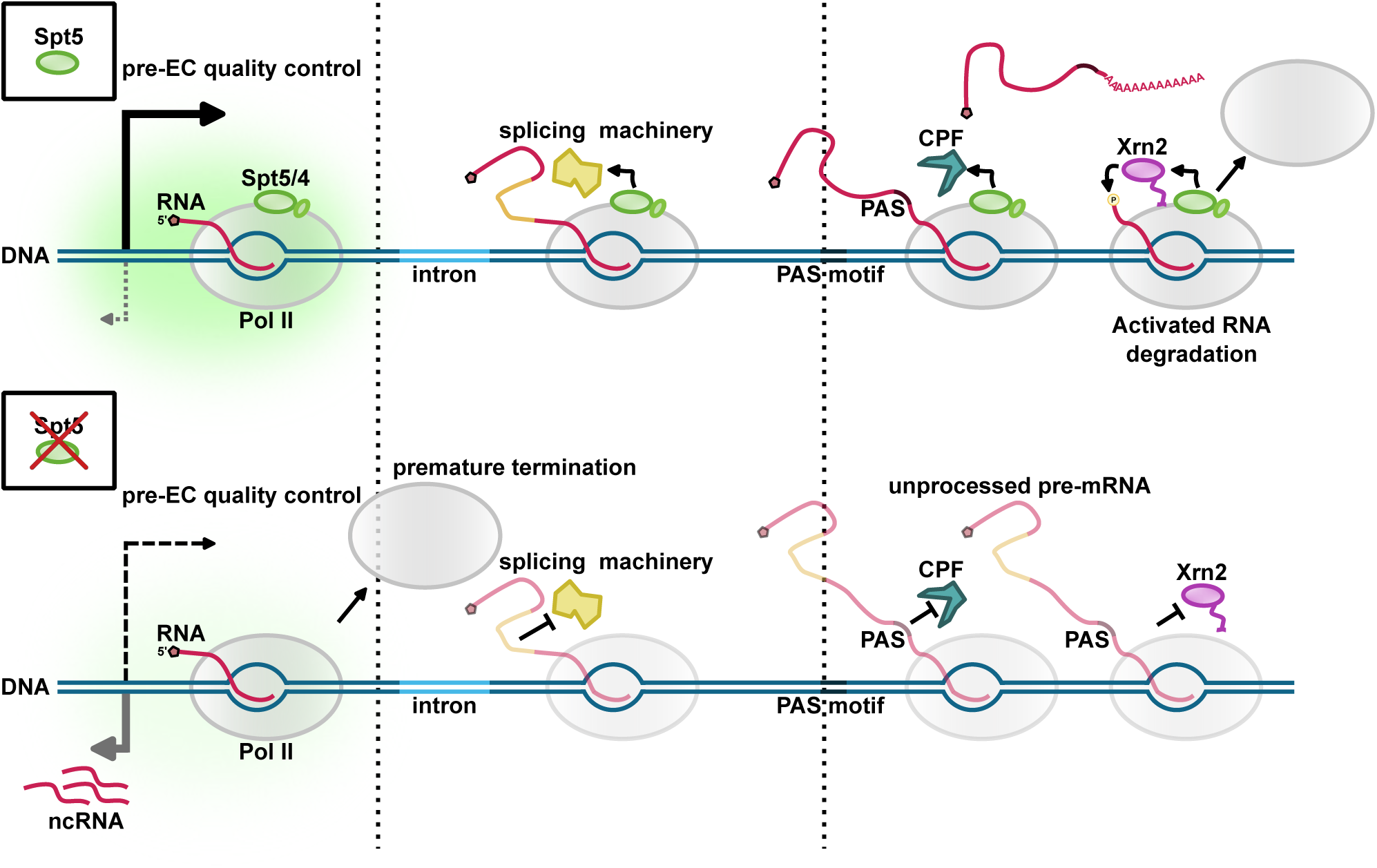
Multifaceted roles of Spt5 in transcription including interplay with Xrn2 transcription termination. Spt5 is required for efficient quality control of pre-elongation Pol II complex (pre-EC) and its absence leads to premature termination on multiple genes. Spt5 plays a role in “licensing” Pol II for efficient co-transcriptional processing (including splicing and 3’end cleavage). Xrn2 uses a short region to anchor itself to the core of Pol II. Spt5 might be stabilising the Xrn2-Pol II complex and provide a context-dependent stimulation of ribonucleolytic activity.

Despite a prominent decrease in transcription for the majority of genes upon depletion of Spt5, several TUs show transcription upregulation (1160 TUs). This group mostly comprises TUs encoding for relatively short (median 770bp compared to 1069bp for full annotation), unstable transcripts such as non-coding RNAs (Figure S5B, S5E) generated from intra/intergenic regions, products of the bi-directional genic promoters transcribed in anti-sense direction (Figure S5F). There is also an increase in housekeeping non-coding RNAs (small nuclear and small nucleolar RNAs) that are stable but targeted by the RNA degradation machinery - RNA exosome for 3’ end trimming/maturation (Figure S5G). Although many non-coding RNAs are low in abundance due to their degradation by the nuclear RNA exosome complex, the synthesis rate of non-coding transcripts is also significantly lower compared to protein-coding mRNAs^123,124^. In mammalian cells, transcription of non-coding RNAs produced from the enhancer regions and bidirectional gene promoters can be suppressed through termination of transcription coupled to RNA degradation^1,20,125–130^. However, the Integrator complex that plays a prominent role in this process in higher eukaryotes is not conserved in fission yeast. We demonstrate that in the absence of the Integrator complex, Spt5 plays a central role in restricting non-coding transcription, possibly representing an ancient mechanism to control transcription within the non-coding genome. Therefore, we propose that Spt5/4 plays a role in restricting transcriptional output from promoters of lowly expressed genes such as non-coding genes that are also supressed by the post-transcriptional mechanisms^109,112,131–134^.

Spt5 also contributes to efficient splicing as we observed a prominent defects for genes with introns in agreement with the published work^84,117,135–139^ suggesting that Pol II complexes escaping promoter-proximal checkpoint are not able to support efficient Pol II elongation and pre-mRNA processing in the cells depleted of Spt5 (Figure S5H, I). We conclude that Spt5 is involved in maturation and quality control ensuring functional coupling of transcription and processing of pre-mRNA.

### Spt5 is required for transcription termination

To explore the genome-wide role for Spt5 in transcription termination, we analysed the TT-seq signal downstream of the annotated PAS for the selected group of non-overlapping TUs analogous to the analyses performed for the Xrn2 depletion (3190 coding TUs). Striking accumulation of the reads downstream of the analysed TUs was observed globally as well as for the individual TUs indicative of a readthrough transcription in Spt5-depleted cells compared to control (Figure 5B, E and F). Although the degree of the readthrough accumulation is milder in Spt5-depleted cells compared to what is observed in Xrn2 catalytic mutant, this is consistent with Xrn2 activity being only partially reduced in the absence of Spt5 (compare Figure S5F to S4F and Figure 3A, F). The role of Spt5 in transcription termination is also supported by the results of the quantitative assay measuring activity of Pho1 in the cells lacking Spt5 (Figure S5J). To rule out that the reduction in Pho1 activity is due to impaired transcription elongation, we also assessed the effect of Spt5 depletion on the activity of Pho1 in a strain lacking promoter driving expression of non-coding RNA. Although in the absence of non-coding transcription Pho1 expression was slightly reduced upon loss of Spt5, repression of Pho1 expression was much more pronounced in the presence of non-coding transcription suggesting that Spt5 plays a role in suppressing *prt* transcription via premature transcription termination.

After Pol II transcribes over PAS, the pre-mRNA is cleaved downstream of the AAUAAA consensus sequence by the CPA complex. Next, we assessed whether pre-mRNA cleavage is affected in Spt5-depleted cells by RT-PCR using a primer pair that spans the PAS region for the representative genes *qcr9* and *spbc21b10.08c*. RT-PCR analyses demonstrated increased readthrough levels across the cleavage site upon Spt5 depletion compared to WT (Figure 5G) suggesting that Spt5 promotes pre-mRNA 3’end cleavage by the CPA. Additionally, when we select PAS sites^140^ and plot read density in a 10bp window normalised to the gene body, we observe a significant shift in distribution (Figure 5H). This might indicate that Pol II complexes that escape Spt5 “licencing” are not able to support pre-mRNA 3’end processing in addition to splicing. Interestingly, depletion of Xrn2 also leads to genome-wide splicing defects and reduced processing efficiency at PAS (Figure 5I, 5G and S5K). This is likely to be due to the extent of transcription interference resulting from the readthrough of upstream genes in the Xrn2 catalytic mutant (Figure S5L), which may prevent proper assembly of functional elongation complex and compromise all steps of pre-mRNA processing.

Together, our data suggest that Spt5 is required for efficient splicing and 3’end processing of pre-mRNA by the CPA machinery as well as for the efficient degradation of the downstream cleavage product by Xrn2 and hence is a central player in coordinating transcription and maturation of pre-mRNA.

## DISCUSSION

In this study, we aimed to identify factors and mechanisms that underlie the function of the essential and conserved transcription factor Spt5. Unexpectedly, we discovered a previously unknown role for Spt5 in coordinating activities involved in RNA processing and transcription. We demonstrated that Spt5 directly interacts with and activates the key transcription termination factor, the RNA 5’-3’ exonuclease Xrn2. Furthermore, we show that loss of Spt5 leads to the accumulation of transcriptionally engaged Pol II at the promoter-proximal region, a severe reduction in transcription elongation/RNA synthesis rate across the gene body and an increase of 3’ readthrough downstream of PAS observed for protein-coding genes. Accumulation of Pol II over 5’ region of TUs in fission yeast was also reported by the previous study that employed NET-seq and ChIP-seq approaches to assess Pol II distribution upon depletion of Spt5^117^. Another study using precision run-on sequencing demonstrated that Spt4, which is non-essential for cellular viability in fission yeast, is also required for promoter proximal pausing and these results are consistent with the role of DSIF (Spt5/4) in the control of Pol II pausing reported in higher eukaryotes^118,141^.

DSIF can stabilize paused Pol II by facilitating the recruitment of NELF^142^. Phosphorylation of Spt5 and NELF by Cdk9/pTEFb leads to the release of Pol II into elongation^60,77,79,143^. However, in organisms such as fission yeast which lack NELF, Pol II pause release is not strictly dependent on Cdk9^72^. Accordingly, depletion of NELF also alleviates dependency on Cdk9 in higher eukaryotes^144^. Additionally, Spt5 can also control pausing independently of NELF^60,145,146^, which might represent a more ancient regulatory mechanism. Indeed, the NGN domain of Spt5 facilitates Pol II pausing by interacting with the non-template strand of DNA, reminiscent of the mechanism described for NusG, the Spt5 homologue in bacteria^62,147,148^.

Our TT-seq data also revealed that upon depletion of Spt5 the fraction of Pol II that escapes into elongation shows defective pre-mRNA processing: splicing and 3’end cleavage at the PAS in addition to altered elongation rate. Reduced splicing efficiency when Spt5 is inactivated or depleted was also observed in other organisms^117,136–138^. These findings are consistent with the idea that Pol II undergoes a “maturation” process during the early stages of transcription and only a small fraction of Pol II is configured for efficient elongation and pre-mRNA processing^149^. At metazoan promoters, a large fraction of transcription initiation events is prematurely terminated^150^ where two endonucleolytic cleavage complexes, CPA and Integrator, cleave nascent RNA to mediate dislodgement of Pol II^20,151–156^. Upon depletion of the Integrator subunits, Pol II can only elongate short distances from the promoter due to low processivity^127,128^ and there is a widespread splicing defect^20^, resembling the profile that we observe upon Spt5 depletion. Our data are consistent with the model where Spt5 contributes to quality control of Pol II complexes during the transition from initiation to elongation stages of transcription. We propose that inefficient splicing and 3’end processing detected in the absence of Spt5 arise due to the leakage of misconfigured Pol II complexes from promoter into gene body/termination zone.

The choice between entry into productive elongation and premature termination is a subject to regulation. Protein phosphatases such as PP1, PP2A and PP4 contribute to premature termination by antagonising Cdk9 activity on Spt5 and Pol II^47,155,157,158^. On the other hand, CPA-induced premature termination can be prevented by the U1 complex of the spliceosome^159,160^. U1 antagonises recruitment of CPA and premature termination by stimulating the Pol II elongation rate at A/T rich regions where Pol II shows low intrinsic processivity in the absence of U1^83,161^. A/T-rich sequences can promote transcription termination by either mediating recruitment of CPA or by destabilising the Pol II complex and making them more susceptible to termination via backtracking-induced pausing^161–164^. Interestingly, we observe that the accumulation of Pol II at the promoter-proximal regions in Spt5-depleted cells correlates with the occurrence of T-rich stretches. This observation is consistent with the recent report demonstrating an increased termination of budding yeast Pol II on poly-U tracks^82^. Spt5 depletion leads to a sharp drop in transcription within the gene body possibly due to destabilisation and premature dissociation of Pol II. We hypothesise that Spt5 controls entry into elongation for properly assembled Pol II complexes by facilitating transcription through T-rich sequences. This group includes TUs encoding for proteins that are subject to regulation in response to various stimuli such as stress or nutrient availability. Future work will be needed to determine how Spt5 is responsible for mediating transcriptional responses to environmental cues.

While depletion of Spt5 leads to decreased transcription elongation across most of the protein-coding genes, transcription of non-coding transcripts is increased. This includes TUs encoding for non-coding transcripts produced from bi-directional genic promoters, and other non-coding transcripts that are intrinsically unstable. Previous studies demonstrated that depletion of the Integrator subunits leads to increased non-coding transcription suggesting that premature transcription termination plays a role in restricting non-coding transcription^20,127,128^. Uncontrolled non-coding transcription when Integrator recruitment is compromised leads to the accumulation of deleterious RNA-DNA hybrids (R-loops) and increased genome instability^165^. Although simple eukaryotes such as yeast lack Integrator, mechanisms are in place that function to restrict and terminate non-coding transcription. In budding yeast, the Nrd1-Nab3-Sen1 (NNS) complex orchestrates the termination at non-coding and short protein coding TUs^19,166^ as well as restricts unwanted non-coding transcription^167^. NNS also mediates the degradation of non-coding RNA by facilitating the recruitment of the RNA exosome complex^168^. Our findings help to explain how fission yeast where the function of the NNS complex is not conserved^31,169^, controls non-coding transcription by instead relying on more universally conserved factor such as Spt5. We propose that restriction of non-coding transcription by Spt5 represents an ancient mechanism that has evolved as a means to defend genomes against the integration of foreign DNA, a role that was demonstrated for a bacterial homologue of Spt5, NusG^170^. Since non-coding transcripts do not usually undergo efficient splicing, Pol II complexes at non-coding promoters may resemble “misconfigured” Pol II and may not be licenced to enter elongation by Spt5. We conclude that Spt5 provides control of maturation and/or licencing of functional Pol II elongation complex at genic promoters to ensure efficient transcription and processing of protein-coding pre-mRNA and at the same time restricts the production of non-coding transcripts in organisms such as fission yeast, which lack the Integrator complex.

Notably, we demonstrate that Spt5 also directly interacts with and enhances the co-transcriptional RNA exonucleolytic activity of Xrn2 to promote efficient transcription termination at the end of the genes. Thus, on functional Pol II complexes that have passed through promoter-proximal quality control and are released into elongation, Spt5 functions to directly modulate the activity of enzymes involved in the RNA processing. Our findings revise the model where Xrn2 chases after transcribing Pol II to engage with the 5’-PO4 ends of the nascent RNA after it is cleaved by CPA^18,19^. It is implied that during 5’ to 3’ degradation of the nascent RNA downstream of PAS, Xrn2 tracks along the nascent transcript until it encounters the RNA polymerase. We provide evidence that Xrn2 forms a stable complex with Pol II which supports the idea of Xrn2 travelling together with elongation complex. This aligns with the observations from other studies as Xrn2 was detected at gene promoters by ChIP^54,85^ and it was implicated in the premature dislodgement of Pol II at metazoan and fission yeast promoters^54,85,171^.

De-phosphorylation of Spt5 CTR by PP1 is required for efficient transcription termination^36,53,78^ which is associated with slowdown of Pol II progression at the 3’end of the TUs^53^. Spt5 stimulates Xrn2-mediated RNA degradation independently of its phosphorylation status. These findings disfavour the idea that de-phosphorylation of Spt5 promotes termination by facilitating Xrn2 engagement with the 5’ end of RNA. Thus, Spt5 de-phosphorylation and deacceleration of Pol II could play a role in the termination of transcription by either contributing to recruitment of CPA, selection of PAS or facilitating conversion of Pol II complex into a “termination competent state”. Indeed, mutants of Pol II that show altered rates and propensity to pause show defective 3’ end processing leading to change in the 3’UTR length and “termination window”^51–53^. However, evidence demonstrating that the dominant negative expression of an Xrn2 mutant can be rescued or exacerbated by Pol II speed mutants strongly suggests that Pol II velocity also contributes to the transcription termination downstream of PAS selection and nascent RNA cleavage by CPA^54^. Further, the cleavage-independent mechanism that relies on the Nrd1-Nab3-Sen1 (NNS) complex mediating termination of non-coding transcripts in budding yeast is also affected by the slow and fast mutants Pol II^55,105^. The altered speed of transcription observed in Pol II mutants can lead to lethality in plants and embryonic lethality in mice. Increased rate of transcription and an altered splicing landscape is associated with ageing further emphasizing the importance of understanding how transcription is coordinated with RNA processing^172–174^. We propose that Spt5 globally orchestrates transcription and RNA maturation as 1) it controls the speed of Pol II in a phosphorylation-dependent manner, 2) it prevents incompetent Pol II complexes from entering elongation and 3) it directly impacts the activity of the RNA processing enzymes recruited to functional polymerases. Specifically, in the context of transcription termination, our data support a model where recognition of the PAS by the CPA triggers Spt5 dephosphorylation by PP1, which slows down Pol II. At the same time, Spt5 directly stimulates Xrn2 activity, assuring highly efficient transcription termination. Our discoveries provide the molecular basis for the previously reported accumulation of the 3’ readthrough upon depletion or small molecule inhibition of Spt5 in mammalian cells^60,85,175^. Xrn2 was also enriched in Spt5 pull-down from mammalian cells suggesting that Spt5-Xrn2 link is conserved^60^. Our cross-linking mass spectrometry data and biochemical experiments demonstrate that Spt5 interacts with Xrn2 and stimulates its activity via its KOW5 and CTR domains. Accordingly, the Spt5 mutant lacking KOW5 and CTR shows a global accumulation of the 3’extended RNA in *S. cerevisiae*^176^.

Recent findings demonstrated that transcription factors that form elongation complex are important for mediating RNA-processing during Pol II transcription^60,177–179^. This includes factors such as components of PAF1 complex, Spt6 and bromodomain protein BRD4. Structural studies demonstrated that in addition to Xrn2, other RNA processing machinery such as the spliceosome^104^, Integrator^180^ as well as CPA^181^ form complexes with Pol II. Interestingly, our cryo-EM structure demonstrates that Xrn2 binds to the same surface of Pol II that as the U1 complex of the spliceosome^104^, which may indicate that there is a competition between RNA-processing complexes during the transcription cycle. Spt5 was shown to stimulate RNA cleavage by the Integrator complex^180^, further supporting the emerging role of Spt5 in coordinating RNA processing and transcription.

The need for regulation of the enzymatic activity may be particularly important in the context of the functional coupling between two processes. For example, in bacteria, where transcription is spatially and temporarily coupled to protein synthesis, NusG, the bacterial homologue of Spt5, interacts and activates RNA helicase activity of the termination factor Rho^86,182,183^. Although in eukaryotes, protein translation is spatially separated from transcription, it is coupled to RNA degradation, where cytoplasmic paralogue of Xrn2, Xrn1 plays a key role in co-translational degradation of mRNAs^184–186^. Although the core of both exoribonucleases is highly conserved, the middle region of these proteins has undergone evolutionary diversification. Xrn1 uses this region to mediate interaction with ribosomes^186^, whereas Xrn2 utilizes it to provide nuclear localisation signal and to anchor enzyme to Pol II (and potentially Spt5). The exact mechanism of how Spt5 and other co-factors regulate activity of the Xrn-like enzymes remains to be understood in the future. We envision that Spt5 could facilitate the engagement of Xrn2 with the RNA substrate or allosterically stimulate the enzymatic activity of the enzyme.

## METHODS

### Strain construction and growth

*S. pombe* strains were grown in YES medium at 30°C to OD_600_ of 0.4 - 0.7 before harvesting. Standard homology-based methodology for genomic integration was used for either epitope tagging, gene deletion or integration. Xrn2 additional copy (rescue) was targeted to *ura4* locus under native promoter. Strains used in this study are listed in Table S6.

### TT-seq and Bioinformatic Analyses

The TT-seq experiments were performed as previously described with modifications^80,81^. Strains were grown in YES media (with reduced uracil to 10mg/L) to OD_600_=∼0.5 and cultures were treated with DMSO (dimethyl sulfoxide) or 1mM auxin for 2h at 25°C. After depletion, cells were labelled with 5mM 4-tU for 6 min, harvested and frozen in liquid nitrogen. Before RNA extraction OD of *S. pombe* was adjusted and mixed with a fraction of 4-tU labelled *S. cerevisiae* (spike-in, 100:1 ratio). RNA was extracted using the hot-phenol method. RNA (100µg) was treated with DNase and fragmented using NaOH, followed by attachment of MTSEA-biotin-XX linker. Biotinylated RNA was purified using µMACS Streptavidin Kit. Specificity of the labelled RNA enrichment was accessed comparing pull-down enrichment of RNA from non-4-tU labelled cells. The size distribution of purified RNA was evaluated by a bioanalyzer. Sequencing libraries were prepared using 50ng RNA according to the manufacturer’s recommendation for the NEBNext Ultra II Directional RNA Library Prep Kit for Illumina (NEB) and sequenced on the Illumina NextSeq 500.

Reads were quality controlled and trimmed using fastp software^187^, aligned to a concatenated genome (*S. pombe* and *S. cerevisiae*) using STAR^188^. Reads were split by species-specific chromosome names and *S. cerevisiae* uniquely mapped reads were obtained using SAMtools^189^ and used for spike-in normalisation. Differential gene expression analysis was carried out using DESeq2^190^ using 2022 version of PomBase annotation^114^. Gene ontology enrichment was performed using a web-based server AnGeLi^191^. Metagenes and heatmaps were prepared with deepTools^192^ (presented as a log ratio and scaled to gene body when indicated) using combined fission yeast genome annotation^113,114^ keeping only coding genes that do not have on the same strand in 250 bp before TSS or after PAS another transcription unit (n=3190).

The intron retention index was calculated as the ratio of intronic reads to surrounding exonic reads using a custom Python script based on PomBase annotation^114^. Selection of genes with premature termination in Spt5 depleted cells was performed using an in-house Python script and genes longer than 600bp were considered. Read density was calculated in 300bp downstream of TSS and divided by read density in the remaining part of the gene requiring at least 1.5-fold change (n=465). Sequence bias in genes with premature termination was evaluated with seqPattern package in R. Potential cleavage defects were estimated as changes in read density spanning 5nt around experimentally generated PAS sites^140^ normalised to 400 bp read density in gene body (100bp from PAS site). PAS sites were filtered to select the most prominent one and closest to the annotated coding transcript end and contain PAS motif (any of: AATAAA, AATGAA, AATAAT, TAATAA, AAATAA, AATAAT, ATAATA, AATAAT) in 50 bp upstream sequence. Data was plotted as scaled frequency.

### Purification of native Spt5 complexes

Flag-tagged Spt5 was purified from WT as well as mutant cells lacking the *dis2* gene or strains containing T1A or T1E mutations in all the repeats of the Spt5 CTR^36^. As a mock control strain without tag on Spt5 was used. Cells were grown to OD 1.5, harvested and cell pellets were frozen at -80°C. Cells were disrupted in a freezer mill (SPEX SamplePrep) in liquid nitrogen. For purification, 5g of cell powder was resuspended in 25ml of lysis buffer (buffer L) (25mM TRIS pH 7, 100mM NaCl, 0.5mM MgCl2, 0.5mM β-Mercaptoethanol) supplemented with 1mM PMSF (phenylmethylsulfonyl fluoride), cOmplete Protease Inhibitor Cocktail (Roche) and phosphatase inhibitors: 1mM NaF, 1mM sodium orthovanadate, 2mM imidazole, 1mM sodium pyrophosphate decahydrate, 0.5mM glycerol-phosphate, 5µM cantharidin, 5nM calyculin A. The lysates were incubated 30 min in the cold-room and cleared at 40000g for 20 min at 4°C. Following centrifugation, the lysates were incubated with 250µl of an equilibrated anti-Flag M2 agarose slurry for 1h in a cold room. Beads were washed 10 times with 1ml of buffer L and proteins were eluted twice with 150µl of 0.2M glycine pH 2.5. Samples were neutralised with 25µl of 1M TRIS. The samples were denatured with 8M urea, reduced with TCEP, alkylated with 2-chloroacetamide, and digested with LysC, and trypsin. The reactions were stopped with formic acid and subjected to analysis by mass spectrometry. Results were analysed using the R environment. For the volcano plot, a protein was considered if present in both replicates of IP with a minimum of two PSM (peptide-spectrum match). Data was median normalised, and missing values in the mock experiment were imputed based on minimum value. Statistics were performed with the DEP package. Data for the mutants was plotted without normalisation as similar numbers of peptides were recovered in each sample and proteins were required to have at least 3 PSM and 3-fold change over mock IP.

### Pol II purification

Large-scale *S. pombe* Pol II purification was performed essentially as in^36^ with the following modifications. Cells were disrupted in a freezer mill (SPEX SamplePrep) or French Press (two passes at 35kpsi). The cell lysate was incubated with anti-Flag M2 beads for 1.5h. Proteins were eluted with 5 ml of 1 mg/ml Flag-peptide (Sigma), followed by the addition of 20 ml of QA buffer (50mM TRIS pH 7.7, 5mM NaCl, 10% glycerol, 0.5mM MgCl_2_, 0.5mM Mg(OAc)_2_, 1mM β-mercaptoethanol). Protein eluate was then applied to an ion exchange chromatography column (2x 1ml HiTrap Q HP, GE Healthcare) equilibrated with QA buffer. The column was washed with several column volumes of 8% buffer QB (same as QA, except 2000mM NaCl) until a stable baseline was achieved. Protein was eluted with a gradient of QB buffer (up to 40%). Pol II buffer was exchanged to CB buffer (25mM HEPES pH 7.9, 100mM NaCl, 1mM MgCl_2_ and 1mM β-mercaptoethanol) and concentrated with Vivaspin 50kDa MWCO (GE Healthcare). Pol II was aliquoted and snap-frozen in liquid nitrogen until the day of the experiment.

### Recombinant protein expression and purification

C-terminally truncated Xrn2 (residues 1-885) His8-tagged^101^ and full-length Rai1 Strep-tagged were expressed from pRSFDuet plasmids in Rosetta *E. coli* strain, grown at 37 °C following induction with 0.3mM IPTG for 12-15h at 20°C. Cells were collected by centrifugation at 4 °C, 5000g for 10 min. For Xrn2, frozen pellets were re-suspended in NA buffer (50 mM Tris-HCl pH 7.5, 500mM NaCl, 5 mM imidazole, 1mM β-mercaptoethanol) supplemented with protein inhibitor cocktail, followed by lysis in French Press and addition of PMSF to 1mM. Lysates were cleared at 4 °C, 40000g for 20 min, filtered and loaded onto a nickel-nitrilotriacetic acid resin and incubated for 30 min. Proteins were eluted with 200mM imidazole. Fractions containing protein were concentrated and separated on HiLoad 16/600 Superdex 200 (Cytiva) equilibrated in CB buffer. Full-length Rai1-Strep was purified on StrepTrap HP (Cytiva) column in CB buffer with 500mM NaCl and eluted with desthiobiotin, followed by gel filtration in CB buffer using Superdex 200 (10/300, Cytiva). Proteins were aliquoted, snap-frozen in liquid nitrogen and stored at -80°C. Full-length Spt5/4 heterodimer (Spt5 containing N-terminal His8-tag and thioredoxin) was expressed from pRSFDuet plasmid and purified as described before^36^ with the exception that ion exchange purification was replaced by Heparin Sepharose (HiTrap Heparin HP Columns, Cytiva) and GF buffer was same as for Xrn2. KOW5-sCTR was expressed either as His-thioredoxin fusion (like Spt5/4 and used for degradation assay using fluorescence anisotropy) or N-terminally His-tagged (used for binding assay and degradation assay). Purification of these constructs was essentially identical to full-length Spt5/4 with the exception that gel filtration was performed on HiLoad 16/600 Superdex 75 (Cytiva).

### Cryo-EM complex preparation

Pol II was mixed with RNA/template DNA at RT for 15 min. After, non-template DNA was added and kept at RT (scaffold 1, Table S2). Spt5 was supplemented (∼3-fold molar excess) and put on ice for 10 min. The complex was mixed with M2 Flag resin (150µl of washed slurry) and incubated in the cold room for 30 min. Beads were washed four times with sample buffer (SB: 25mM HEPES pH 7.9, 100mM NaCl, 0.5mM MgCl_2_, 0.05% Tween) and incubated on ice for 30 min with an excess of Xrn2^M^/Rai1. The resin was washed 5x with 150µl SB buffer and complex eluted two times with 35µl of 2.5mg/ml 3xFlag peptide (10 min incubation each). Sample buffer was exchanged against SB buffer using a 50kDa amicon device. The complex was crosslinked with 0.1% glutaraldehyde for 10 min on ice and quenched with 8mM aspartate, and 2mM lysine. Finally, the complex buffer was exchanged for the SB buffer as before. Grids were prepared using a Vitrobot mark IV (Thermo Fisher Scientific) at 100% relative humidity. Quantifoil Holey Carbon R2/1 200 mesh gold grids were glow discharged, before applying 3.5µL of sample at around 0.7 mg/ml and blotted for 3.5s, blot force -25 before vitrification in liquid ethane.

### Cryo-EM image collection and processing

Cryo-EM data were collected at the Oxford Particle Imaging Centre (OPIC), on a 300kV G3i Titan Krios microscope (Thermo Fisher Scientific) fitted with a K2 Summit (Gatan) direct electron detector and a GIF Quantum energy filter (Gatan). Automated data collection was setup in SerialEM v3.8 and movies were recorded in counting mode. Data were collected using a 3×3 multishots pattern, with a total dose of ∼ 42.5e^-^/Å^2^, split across 50 frames, a calibrated pixel size of 1.05 Å/px and a 20eV slit. Sample-specific data collection parameters are summarised in Table S4. Data were processed using the cryoSPARC V-3.X^193^, following standard workflow. Pre-processing was performed using patch motion correction and patch-CTF estimation with default settings. Corrected micrographs with poor statistics where manually curated. A first round of blob picking followed by a round of 2D classification allowed to generate initial templates that were used for template picking. After 2D classification, the high-resolution classes were selected, and five *ab-initio* models were generated and further refined using heterogeneous refinement. Only, the class containing the DNA duplex and RNA was selected and the 984,000 particles were refined using NU-refinement. After refinement of the CTF and correction of second order aberration per optics group, the particles were refined a second time with NU-refinement before being exported to RELION 3.1^194^ for further 3D classification. Based on crosslinking mass spectrometry results, we designed a mask surrounding most of the Pol II residues involved in these crosslinks and performed a 3D classification without alignment, using 10 classes and T=4. After visual inspection, one class containing 215000 particles had an extra density for both the Kow5 domain of Spt5 and a fragment of Xrn2. The particles belonging to that class were re-imported into cryoSPARC and a final NU-local refinement was performed, leading to a 2.67 Å resolution map.

### Structure determination and model refinement

Each of the individual subunits were first modelled by first fitting the individual domains predicted by Alphafold2 implemented in CollabFold^103,195^. In WinCoot^196^, the restraints module was used to generate restraints at 4.3Å and allow flexible refinement to fit the main chain into density. Multiple cycles of manual adjustment in WinCoot followed by real refinement in PHENIX^197^ were used to improve model geometry. The final model geometry and map-to-model comparison was validated using PHENIX MolProbity^198^. All map and model statistics are detailed in Table S4. Structural analysis and figures were prepared using UCSF ChimeraX^199^.

### RNA degradation activity assays/RNA binding

Pol II was incubated with an excess of annealed RNA/template-DNA at RT and mixed with non-template DNA (scaffold 1a with P-RNA-FAM, Table S2). This complex was mixed with equilibrated streptavidin beads (Streptavidin Sepharose High Performance, Cytiva) and incubated at room temperature for 30 min. Beads were washed with TB buffer (20mM TRIS, 40mM KCl, pH adjusted to 7.9 with HCl), W500 (20mM TRIS, 500mM NaCl pH adjusted to 7.9 with HCl) and again 3 times with TB buffer. Immobilised Pol II complex (20µl) was mixed with either 10µl SB buffer (25mM HEPES pH 7.9, 100mM NaCl, 1mM MgCl2, 1mM β-mercaptoethanol) or Spt5/4 in SB buffer and incubated for 10 min at RT. Beads were washed two times with TB buffer. The complex was mixed with buffer or Xrn2 and samples were collected after 3 and 6 min of incubation. Samples were resolved on 10% 8M UREA-PAGE to evaluate RNA degradation and visualised for FAM fluorescence.

RNA degradation assays were also performed on RNA substrate (without Pol II). The reaction was set up in RB buffer (10 mM HEPES, pH 7.9, 100mM NaCl, 1mM MgCl2, and 1mM β-mercaptoethanol) using P-RNA-FAM (Figure S1D, Table S2). Aliquots were taken at 5, 10, 20 min and reactions were terminated by the addition of stop buffer (10mM EDTA, 3.5M urea, 50μg/mL heparin, 0.01% bromophenol blue, 0.015% xylencyanol in formamide) and resolved by 10% 8M UREA-PAGE. Alternatively, the kinetics of RNA degradation were studied using fluorescence anisotropy (FA) assay. To this end, degradation P-RNA-FAM (∼250nM) with protein (2-3-fold excess over RNA) or RB buffer was started by the addition of equal amounts of Xrn2 (optimised for the given reaction). Two constructs for KOW5-sCTR were tested without or with thioredoxin (Figure 3E and S3F, respectively). P-RNA-FAM excitation was induced with linearly polarized light at 485 nm and emission was measured both parallel and perpendicular to the plane at 520 nm at 25 °C using a PHERAstar FS plate reader (BMG Labtech). Polarisation anisotropy data were rescaled to 0-1 and analysed with in-house R script and fitted with an exponential curve. KOW5-sCTR (without thioredoxin) binding to 50nM P-RNA-FAM was evaluated using fluorescence anisotropy assay with increasing amounts of protein. The binding curve was fitted as described previously^200^ using the nonlinear least squares procedure in R.

### Termination assay

Pol II was incubated with an excess of annealed RNA/template-DNA at RT and mixed with non-template DNA (scaffold 1a, Table S2). This complex was mixed with equilibrated streptavidin beads (Streptavidin Sepharose High Performance, Cytiva) and incubated at room temperature for 20 min. Beads were washed with TB buffer (20mM TRIS, 40mM KCl, pH adjusted to 7.9 with HCl), W500 (20mM TRIS, 500mM NaCl pH adjusted to 7.9 with HCl) and again 3 times with TB buffer. Immobilised Pol II complex (20µl) was mixed with either 10µl reaction buffer (RB) (25mM HEPES 7.9, 100mM NaCl, 5mM MgCl_2_, 4mM ATP) or proteins in RB. Reactions were incubated for 10 min at 25°C and supplemented with an additional 300mM NaCl. Beads were spun down, and the supernatant was filtered through the Costar-X column to remove any potential beads. The remaining beads were washed with TB buffer. The reaction was inhibited with stop buffer (10mM EDTA, 3.5M urea, 50μg/mL heparin, 0.01% bromophenol blue, 0.015% xylencyanol in formamide). Samples were used for Western blotting or UREA-PAGE to evaluate RNA degradation.

### *In vitro* transcription

Pol II complexes were prepared in a similar manner as for termination assay using scaffold 2 with ^32^P-RNA (Table S2). After washes with TB buffer, complex was incubated with buffer or Spt5/4. Transcription was started with the addition of 50µM rNTPs and samples were collected at times specified. Products were resolved on 10% UREA-PAGE gel and visualised on FLA-7000 phosphoimager (Fujifilm).

### Immunofluorescence

Immunofluorescence was performed as previously described with minor changes^201^. Briefly, cells were grown in YES media to a final 0.5OD and 9ml culture was fixed with 1ml 37% formaldehyde for 25 min at room temperature. Cells were washed twice with PEM (100mM PIPES, 1mM EGTA, 1mM MgSO_4_; pH 6.9) and resuspended in PEMS (PEM with 1.2M sorbitol). The cell wall was digested with 0.25mg/ml of zymolyase (100T) at 37°C for 1 hour. Cells were permeabilised with PEMS+1% Triton X-100 for 10 min and washed three times with PEM. Cells were then resuspended in PEMBAL (1% Bovine serum albumin, 100mM Lysine hydrochloride, 0.1% NaN3) and incubated with primary antibody (monoclonal α-V5-Tag mouse antibody /Elabscience/ and monoclonal α-Flag M2 mouse antibody /Sigma Aldrich/) and secondary antibody (AF488-conjugated goat anti-mouse antibody /Elabscience/). Cells in PBS were mounted on poly-lysine-treated coverslips (18mmx18mm, 170±5µm high precision) using DAPI-containing mounting medium (ROTI®Mount FluorCare DAPI) and images acquired on the Deltavision Ultra High-Resolution Microscope.

### RT-PCR

DNase-treated RNA isolated from indicated strains was reverse transcribed with either Superscript III or IV using primers listed in Table S2 (three or one reverse primers were used for cDNA synthesis for gene body (gb) or cleavage site (cs)/termination zone (term) evaluation, respectively). PCR products were amplified using Phire II polymerase at 60°C with following cycles: for the *qcr9* gene: gb fragments were amplified with 30 cycles, cs and term fragments with 28 for Xrn2 strains or 32 for Spt5 degron, whereas for *spbc21b10.08c*: 30 cycles were used for gb and 29 cycles for cs and term fragments.

### Phosphatase reporter assay

Cells were grown in YES media and transferred to EMM-P (no-phosphate media, NaOAc 1.2g/L, L-Glutamic acid monosodium salt hydrate 5g/L, glucose 20g/L, mineral stock, vitamin stock, salt stock as for EMMG^202^, adenine, histidine, uracil, leucine, lysine 225mg/L). OD of cell cultures was adjusted, and cells were treated with DMSO or 1mM auxin for 5h at 25°C. Cells were washed and resuspended in water. Phosphatase activity was measured as the ability of cells to dephosphorylate para-nitrophenylphosphate at room temperature for 10 min (the reaction was stopped with saturated Na_2_CO_3_ and measured at 400nm). Phosphatase activity was normalised to OD and for Spt5 degron it was scaled to the DMSO-treated controls.

### Reconstitution of Pol II complexes and cross-linking

Complex formation between Xrn2^M^, Rai1, Spt5, Spt4 and Pol II with nucleic acid scaffold (Table S2, scaffold 1) was analysed by the size-exclusion chromatography (Superose 6 3.2/300), and Coomassie-stained PAGE.

For crosslinking, Pol II was incubated with ∼2 molar excess of RNA/template-DNA at RT and mixed with non-template DNA (∼3 molar excess, three different scaffolds were tested 1, 1b, 2, Table S2). Excess of Spt5/4 and Xrn2^M^/Rai1 were added. Complexes which were purified as described for cryo-EM or gel filtration were subjected to BS3 cross-linking on ice for 1h. The reaction was quenched with TRIS pH 7.5. Buffer was exchanged using an Amicon concentrator.

Alternatively, the complex was assembled and crosslinked and then purified by Superose 6 3.2/300. Samples were denatured with 8M urea, reduced with TCEP, alkylated with 2-chloroacetamide and digested with LysC, and trypsin. The reaction was stopped with formic acid and subjected to analysis by mass spectrometry.

### Analysis of pull-down or cross-linked peptides by mass spectrometry

Obtained peptides were separated by nano-flow reversed-phase liquid chromatography coupled to a Q Exactive Hybrid Quadrupole-Orbitrap mass spectrometer (Thermo Fisher Scientific). Raw data files were processed for protein identification using MaxQuant, version 1.5.0.35, integrated with the Andromeda search engine as described previously. The MS/MS spectra were searched against *S. pombe* Uniprot database, precursor mass tolerance was set to 20 ppm and MS/MS tolerance to 0.05 Da. Enzyme specificity was set to trypsin with a maximum of two missed cleavages. Protein and peptide spectral match false discovery rate was set at 0.01. For cross-linked peptides raw files were converted into mgf format using pParse and searched using the pLink software^203^. Data was analysed and visualized in UCSF Chimera or using a custom R-script.

### DATA AND CODE AVAILABILITY

The model and map for the structure have been deposited in the PDB: 8QSZ and EMDB: EMD-18643. TT-seq data have been deposited in Gene Expression Omnibus: GSE244546.

### SUPPLEMENTAL INFORMATION

Supplemental information is provided in attached spreadsheet file.

## ACKNOWLEDGEMENTS

We thank Nikolay Zenkin and Soren Nielsen for their help and advice on setting up an *in vitro* transcription system; Tomo Sugiyama and Japanese National BioResource Project (NBRP)^6–9^ for providing *S. pombe* strains for AID-tagging, members of the Vasilieva lab and Aleksander Szczurek for helpful discussions, advice, and valuable comments on the manuscript. This work was supported by the Wellcome Trust Senior Fellowship to LV (WT106994/Z/15/Z), the Wellcome Trust Investigator Award to JMG (200835/Z/16/Z and 222510/Z/21/Z) and the Emmy Noether Programme of the Deutsche Forschungsgemeinschaft (DFG) to CK (KI 1657/2-1). SSH is funded by a scholarship grant (MM1/23/ID:1563097) from the Egyptian Mission Department. Access to electron microscopes was provided by the OPIC Electron Microscopy Facility funded by Wellcome JIF (060208/Z/00/Z) and equipment (093305/Z/10/Z) grants. Access to computational resources was supported by the Wellcome Trust Core Award Grant Number 203141/Z/16/Z with additional support from the NIHR Oxford BRC. The views expressed are those of the author(s) and not necessarily those of the NHS, the NIHR, or the Department of Health.

## AUTHOR CONTRIBUTIONS

K.K. and L.V. conceived, designed the experiments, and wrote the manuscript. K.K. generated strains and plasmids for this study, performed protein purifications, *in vitro* RNA degradation and *in vitro* transcription elongation and termination experiments, sample crosslinking for mass spectrometry experiments, RNA extraction, TT-seq library preparation and the bioinformatics analysis, participated in modelling of cryo-EM structure and mass spectrometry data analysis. E.A. and C.K. performed microscopy experiments to analyse the cellular localisation of Xrn2 mutants and quantification of the results. L.C. and J.M.G. performed cryo-EM experiments including grid preparation, data acquisition and assisted with data analyses. T.K. generated Cdk9/cyclin T1 (Pch1) baculovirus construct and purified Cdk9/cyclin T1 (Pch1) complex from insect cells. M.F. performed mass spectrometry experiments and assisted with analysis. S.S.H. helped with RT-PCR analysis. All authors edited the manuscript.

---

**Figure S1.**
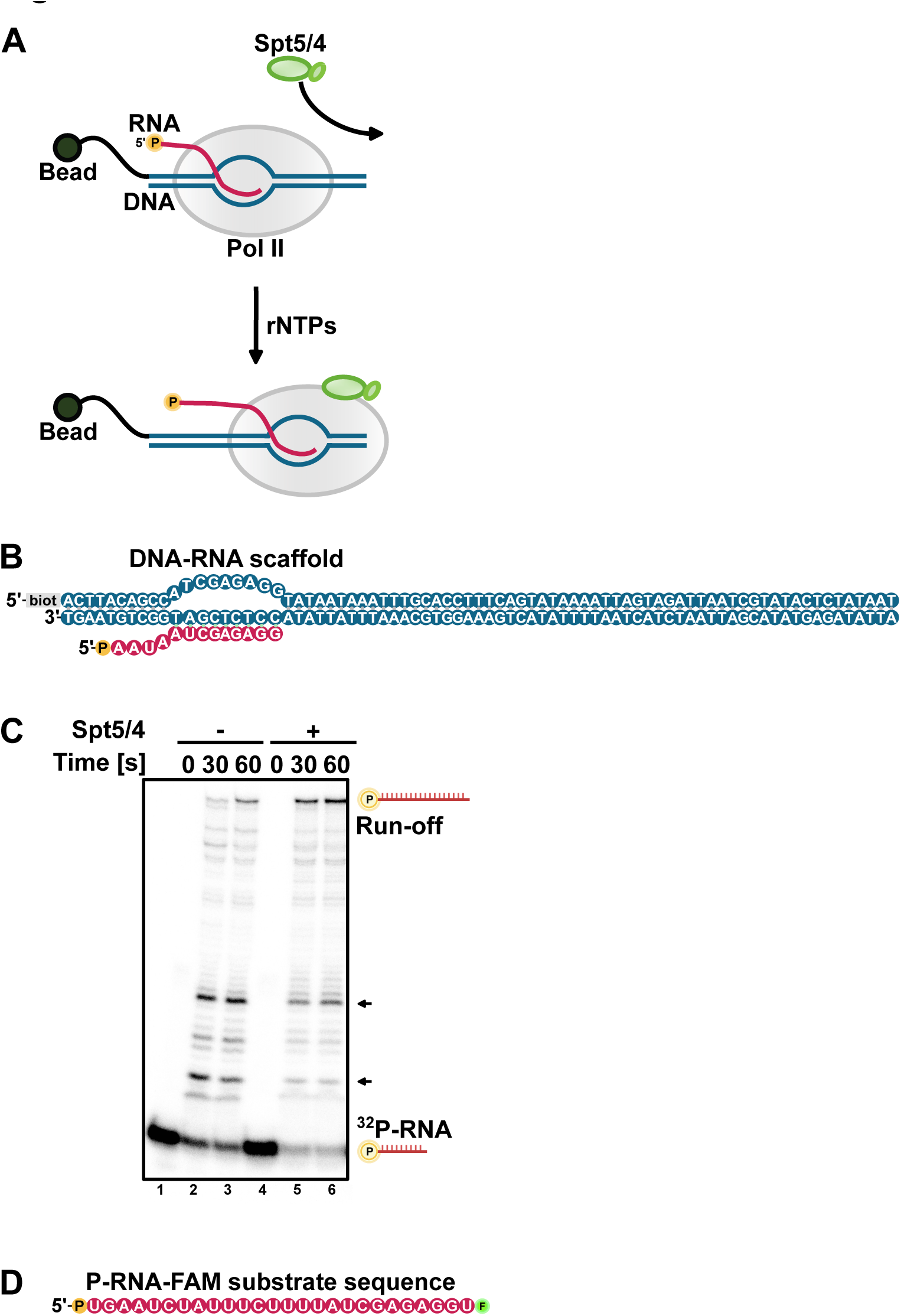
Pol II complex is transcriptionally competent. (A) Schematics of *in vitro* transcription assay. Pol II is assembled with RNA-DNA hybrid and immobilised on beads via biotin on non-template DNA strand (scaffold 2, Table S2). The complex was prepared with or without Spt5/4. Transcription was initiated with rNTPs, and radioactive products separated on UREA-PAGE gel. (B) DNA-RNA scaffold used for *in vitro* transcription (C) Spt5 has a stimulatory effect on transcription *in vitro.* Complexes with or without Spt5/4 were prepared with RNA 5’-labelled with ^32^P and transcription was started with rNTPs. Samples were collected at indicated time points and resolved on UREA-PAGE gel. Arrows highlight lower pausing in the presence of Spt5/4. (D) Schematics of 5’-monophosphate RNA with 3’-FAM fluorescent label (P-RNA-FAM) used for degradation assays.

**Figure S2.**
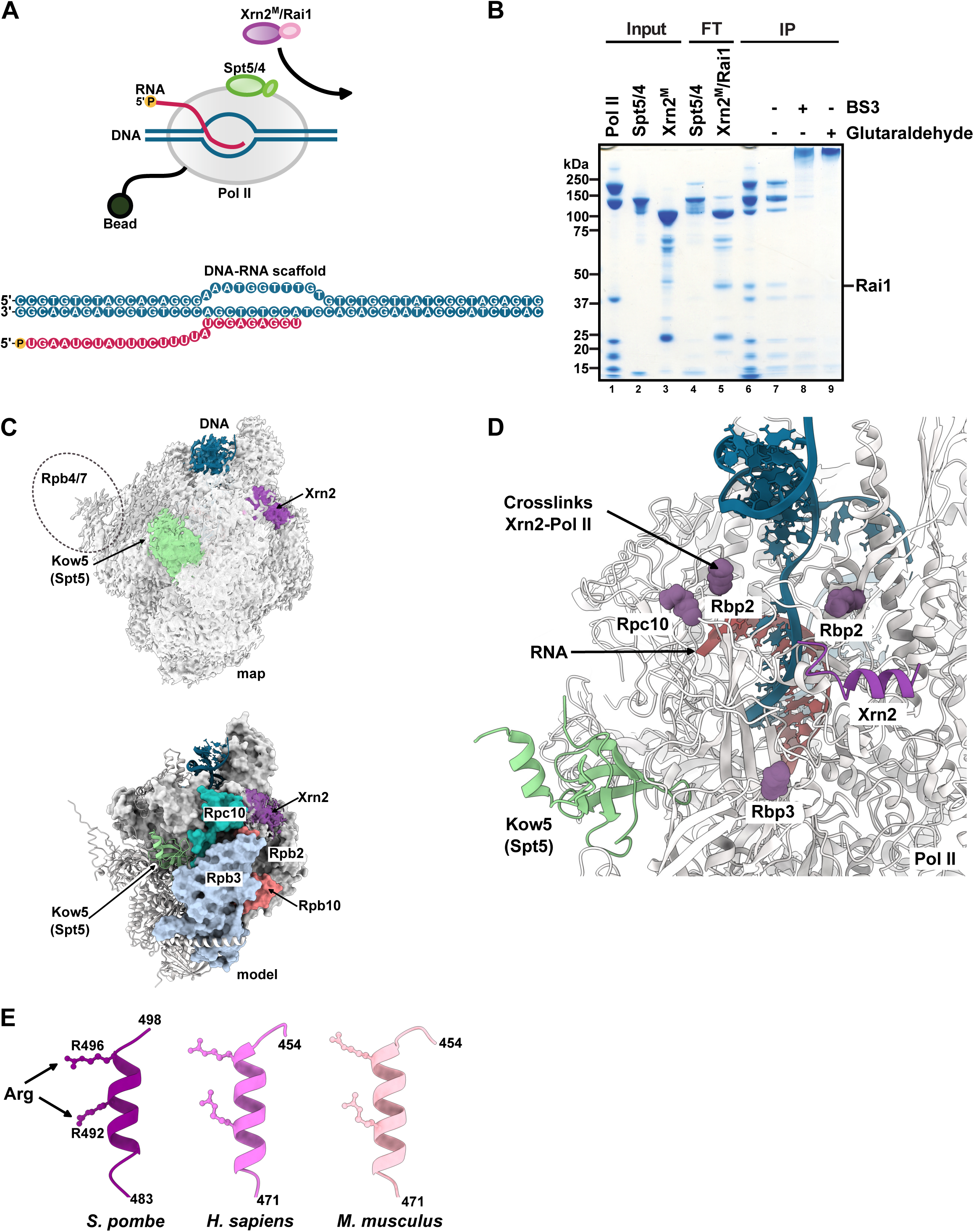
Probing Xrn2 interactions with Pol II. (A) Schematics of *in vitro* Pol II complex reconstitutions. Pol II was immobilised on anti-Flag beads (via Rpb9) and proteins were added in a stepwise manner. Protein excess was washed away, and complexes were eluted with Flag-peptide. The nucleic acids scaffold used is shown in the bottom (B) Xrn2-Rai1-Spt5/4-Pol II-DNA-RNA complex crosslinked with BS3 or glutaraldehyde was analysed by SDS-PAGE gel. FT – refers to the flow-through, IP-immunoprecipitation (C) Depiction of the cryo-EM density for the glutaraldehyde crosslinked complex. Areas of Spt5 KOW5 and Xrn2 peptide binding are highlighted in green and violet, respectively. In the reconstitutions, Rpb4 and Rpb7 were not visible, and their location is marked as an oval. The lower panel depicts the model with Rpb2/Rpb3/Rpc10 shown as surfaces (D) Mapping of the crosslinks between Xrn2 and Pol II on the Pol II/Xrn2/Spt5 structure (compare Figure 2C) (E) Alpha fold prediction of the potential helices that might anchor Xrn2 to Pol II in human and mouse. Two characteristic arginines are shown in the ball-stick representation.

**Figure S3.**
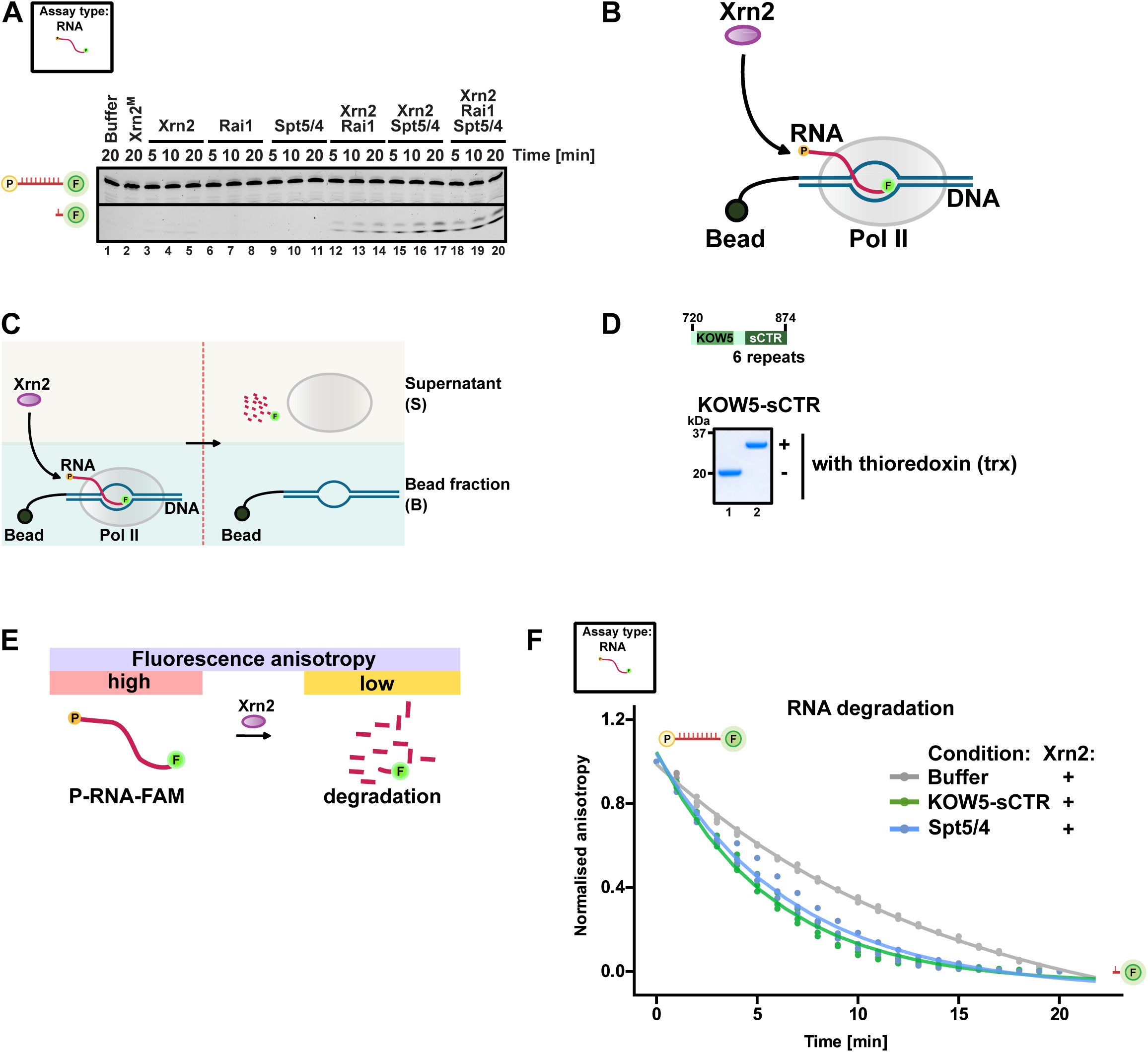
Spt5 affects Xrn2 ribonucleolytic activity. (A) Rai1 and Spt5 can stimulate Xrn2 exoribonucleolytic activity towards 3’FAM labelled 5’-monophosphate-RNA substrate (compare Figure S1D). (B) The schematic diagram for degradation assay with the assembled Pol II complex (scaffold 1a, Table S2). (C) Schematic diagram summarising the *in vitro* strategy for transcription termination assay. Pol II complex is assembled with nucleic acids and immobilised on streptavidin beads using biotinylated non-template DNA. Xrn2**-**mediated transcription termination can be evaluated comparing by supernatant or bead (bound fraction). (D) Purifications of the proteins KOW5-sCTR (either as thioredoxin fusions or His-tag variant protein, compare Figure 3E, F and S3F). (E) Schematics depicting degradation assay based on fluorescence anisotropy. Xrn2 degrades fluorescently labelled RNA, which can be monitored over time. (F) Spt5 stimulates degradation by Xrn2 via fragment containing the KOW5 domain (KOW5-sCTR-Spt5 region 720 to 874 aa). Fluorescence polarisation anisotropy assay comparing RNA degradation kinetics of Xrn2 alone, KOW5-sCTR (compare to Figure 3F) or full-length Spt5/4. Constructs used with N-terminal thioredoxin fusion.

**Figure S4.**
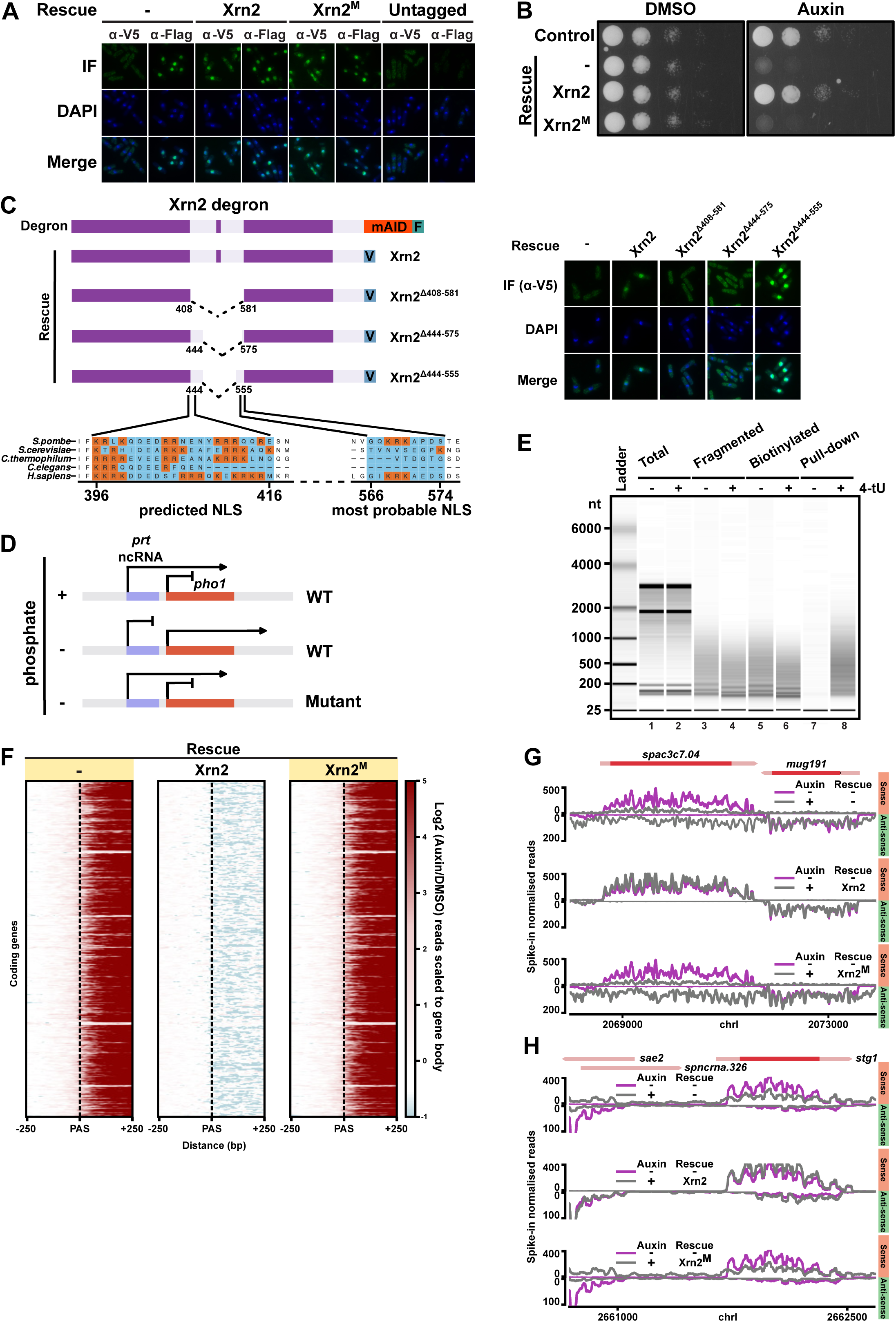
Functional analyses of Xrn2 depleted or mutated strains. (A) Subcellular localisation of Xrn2 mutants. Immunofluorescence (IF), DAPI and merged images are shown. Untagged strain is used to validate the specificity of the antibodies. (B) Growth analysis of strains used in the study grown in the presence or absence of auxin. Expression of two Xrn2 copies does not affect the growth of the *S. pombe.* Cells lose their viability after Xrn2 depletion and Xrn2^M^ cannot rescue cell growth defects. (C) Mapping of Xrn2 nuclear localisation signal. Localisation of different truncation mutants was assessed by immunofluorescence. Predicted NLS is not sufficient to target protein to the nucleus. The most probable NLS is in region 555-575 aa Xrn2 (D) Schematic diagram explaining intrinsic phosphatase reporter assay. In phosphate-rich media, *pho1* gene expression is repressed due to transcription of long non-coding RNA *prt* that interferes with Pho1 transcription. In low phosphate conditions, RNA is terminated prematurely to allow expression of the phosphatase (Pho1). If termination is defective (i.e. upon Xrn2 depletion), *prt* transcription will interfere with *pho1* expression resulting in low phosphatase activity. (E) Evaluation of TT-seq workflow. RNA is only efficiently pulled down if cells were labelled with 4-thiouracil (4-tU) before RNA pull-down. (F) Genome-wide readthrough of coding genes. The region around PAS is presented as a heatmap with a log_2_ fold ratio of the indicated treatment to control strain (DMSO treated strain without Xrn2 complementation). Read density is normalised to the gene body (compare to Figure 5F). (G) Example genome TT-seq tracks to illustrate gene downregulation by readthrough from the opposite strand (compare Figure 4D, cluster 3 and 1). (H) Locus with genes in tandem orientation showing downregulation of the gene by the upstream readthrough transcription.

**Figure S5.**
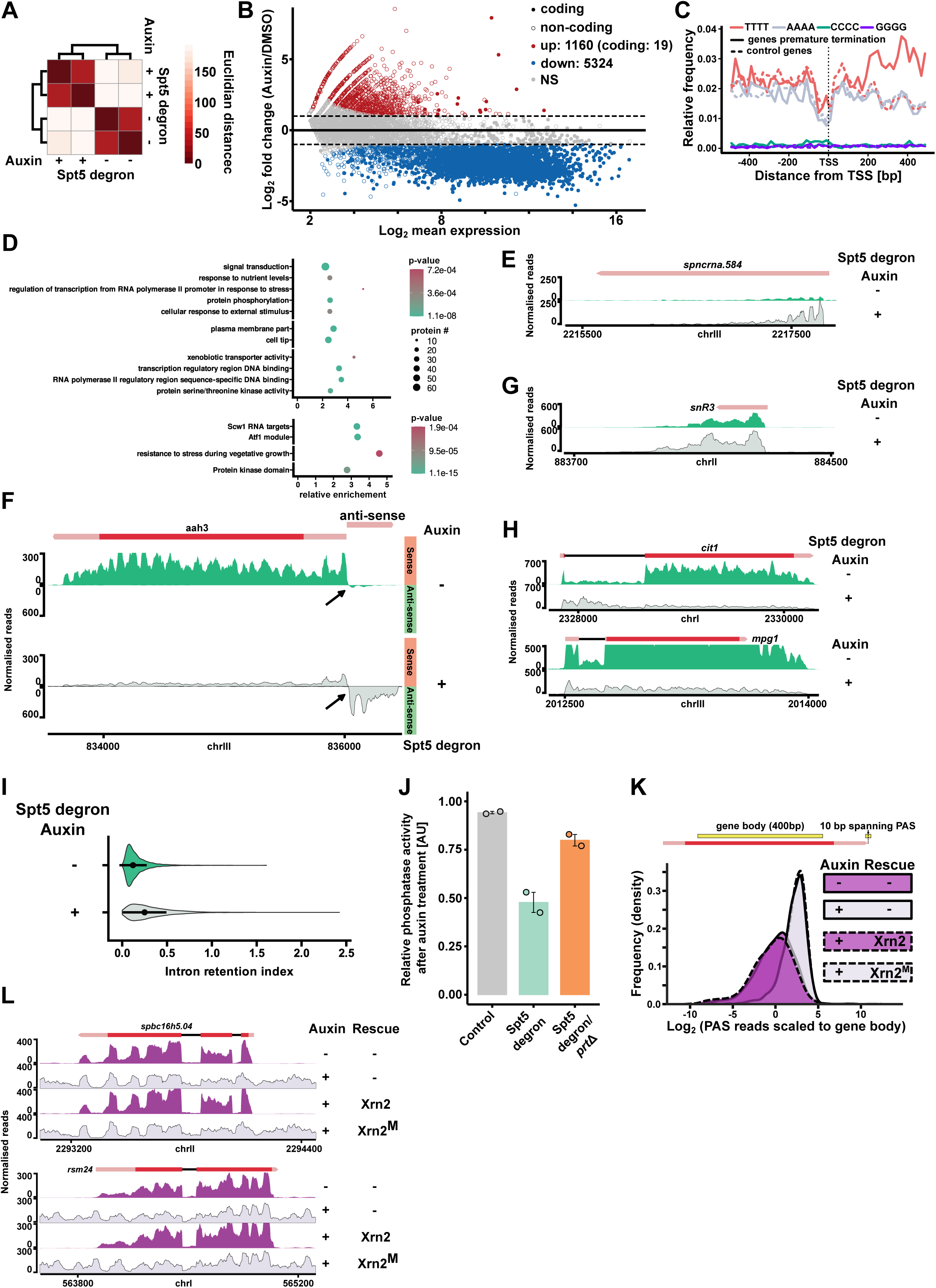
Spt5 loss leads to global transcriptional dysregulation. (A) Heatmap showing Euclidean distances between TT-seq replicates for Spt5 degron (+/-auxin). (B) MA plot illustrating changes in transcriptome upon loss of Spt5. Spt5 depletion leads to global transcription downregulation. Upregulated genes predominantly belong to the non-coding class. (C) Coding transcription units that show premature termination/attenuation (n=465, compare Figure 5D) exhibit higher T content in the TSS region. Runs of nucleotides (at least 4) are shown as a metagene plot. (D) Enrichment gene analysis of coding transcription units that show premature termination/attenuation (n=465, compare Figure 5D). A subset of non-redundant categories included, for the full set refer to Table S5 (E) Depletion of Spt5 upregulates non-coding RNA, an example genomic snapshots (F) Spt5 depletion increases promoter bi-directional anti-sense transcription. (G) Stable non-coding RNA are affected after Spt5 loss. (H) Spt5 impacts splicing, an example genome browser snapshots (I) Violin plots showing intron retention index before and after Spt5 loss highlighting genome-wide splicing defects. (J) Assessment of transcription termination using internal reporter that modulates phosphatase homeostasis (compare Figure S4D). Phosphatase activity serves as a proxy for termination efficiency. Loss of Spt5 correlates with decreased termination efficiency. Strain, where the non-coding RNA (*prt* lnRNA) is deleted, serves as a reference for elongation defects after Spt5 loss. (K) Xrn2 and its catalytic activity might contribute to 3’end processing (analysis as in Figure 5H). (L) Readthrough generated after loss of Xrn2 activity is not properly spliced.

